# A specialized MreB-dependent complex mediates the formation of stalk-specific peptidoglycan in Caulobacter crescentus

**DOI:** 10.1101/389114

**Authors:** Maria Billini, Jacob Biboy, Juliane Kühn, Waldemar Vollmer, Martin Thanbichler

## Abstract

Many bacteria have complex cell shapes, but the mechanisms producing their distinctive morphologies are still poorly understood. *Caulobacter crescentus*, for instance, exhibits a stalk-like extension that carries an adhesive holdfast mediating surface attachment. This structure forms through zonal peptidoglycan biosynthesis at the old cell pole and elongates extensively under phosphate-limiting conditions. We analyzed the composition of cell body and stalk peptidoglycan and identified significant differences in the nature and proportion of peptide crosslinks, indicating that the stalk represents a distinct subcellular domain with specific mechanical properties. To identify factors that participate in stalk formation, we systematically inactivated and localized predicted components of the cell wall biosynthetic machinery of *C. crescentus*. Our results show that the biosynthesis of stalk peptidoglycan involves a dedicated peptidoglycan biosynthetic complex that combines specific components of the divisome and elongasome, suggesting that the repurposing of pre-existing machinery provides a straightforward means to evolve new morphological traits.

## Introduction

The shape of most bacteria is determined by a cell wall made of peptidoglycan (PG), a mesh-like hetero-polymer that surrounds the cytoplasmic membrane and provides resistance against the internal osmotic pressure (Typas et al., 2012; Vollmer et al., 2008). The backbone of PG is formed by strands of alternating *N*-acetylglucosamine (GlcNAc) and *N*-acetylmuramic acid (MurNAc) subunits. These glycan chains are connected by short peptides that are attached to the MurNAc moieties, giving rise to a single elastic macromolecule known as the PG sacculus (Schleifer and Kandler, 1972).

The PG meshwork needs to be continuously remodeled to allow for cell growth and division (den Blaauwen et al, 2008). This task is achieved by a large and highly redundant set of PG synthesizing and degrading enzymes. Insertion of new cell wall material is initiated by the translocation of lipid-linked GlcNAc-MurNAc-pentapeptide precursors across the cytoplasmic membrane to the periplasm (Mohammadi et al., 2011; Sham et al., 2014; van Heijenoort, 2007). Glycosyltransferases (GTases) then incorporate the disaccharide units into preexisting glycan strands, while the L-Ala–D-Glu–L-Lys/*meso*-DAP–D-Ala–D-Ala pentapeptides of adjacent glycan strands are crosslinked by transpeptidases (TPases) (Typas et al., 2012; Vollmer & Bertsche, 2008). Depending on their domain structure, PG synthases can be classified as bifunctional GTases/TPases (class A PBPs), monofunctional TPases (class B PBPs) and monofunctional GTases (Vollmer and Bertsche, 2008). The majority of TPases are DD-TPases, also known as penicillin-binding proteins (PBPs) (Suginaka et al., 1972). These proteins catalyze the formation of D-Ala^4^-*meso*-DAP^3^ (4-3) crosslinks, in a reaction that releases the D-Ala^5^ moiety of the donor molecule (Glauner et al., 1988). Alternatively, crosslinks can also be formed between two *meso*-DAP^3^ residues (3-3 crosslinks), catalyzed by specific LD-TPases that use tetrapeptide side chains as donor moieties and release their terminal D-Ala^4^ residue to gain energy for the crosslinking reaction (Magnet et al., 2008). To remodel the PG sacculus during growth and division, cells require not only synthetic but also lytic enzymes that cleave bonds in the PG meshwork and thus make space for the insertion of new material (Höltje, 1995). Depending on their cleavage specificity, these so-called autolysins can be typically sorted into three main categories. Lytic transglycosylases act on the glycan strands and cleave the β-1,4-glycosidic bond between MurNAc and GlcNAc, leaving 1,6-anhydro-MurNAc as the terminal residue (Scheurwater et al., 2008). Amidases, by contrast, hydrolyze the amide bond between the peptide and the MurNAc moiety (Höltje, 1995), whereas endo- and carboxypeptidases hydrolyze specific amide bonds within the peptides (van Heijenoort, 2011; Vollmer and Bertsche, 2008).

The formation and degradation of PG need to be closely coordinated to prevent cell lysis (Rice and Bayles, 2008; Typas et al., 2012), a task that is presumably achieved by the assembly of synthetic and lytic enzymes into dynamic multi-protein complexes (Höltje, 1998). In the majority of rod-shaped bacteria, two of these complexes have been identified to date. The first one, called the elongasome, mediates the dispersed incorporation of new PG along the lateral walls of the cell during the elongation phase. Its positioning is controlled by the actin-like protein MreB (Daniel and Errington, 2003; Jones et al., 2001; van den Ent et al., 2001), which forms patch- or arc-like filaments that are attached to the inner face of the cytoplasmic membrane (Dominguez-Escobar et al., 2011; Garner et al., 2011; Olshausen et al., 2013; Salje et al., 2011; van Teeffelen et al., 2011). These structures move around the circumference of the cell and, thus, ensure even growth of the rod-shaped sacculus. Their effect on the PG biosynthetic machinery is mediated by the transmembrane protein RodZ (Alyahya et al., 2009; Bendezu et al., 2009; Shiomi et al., 2008), which links MreB to a periplasmic complex containing the elongation-specific monofunctional TPase PBP2 (Lee et al., 2014; Morgenstein et al., 2015; Typas et al., 2012). Towards the end of the elongation phase, PG synthesis is taken over by a second complex, called the divisome (Du and Lutkenhaus, 2017; Typas et al., 2012), which mediates pre-septal elongation and subsequent constriction of the PG sacculus at midcell. Its positioning and activity are regulated by FtsZ, a tubulin homolog that assembles into a dynamic ring-like structure at the future division site. This so-called Z-ring then recruits, directly or indirectly, all other components of the cell division machinery. The divisome includes a variety of PG synthases and hydrolases, among them the division-specific monofunctional TPase PBP3 (Weiss et al., 1999), which act together to coordinately remodel the PG layer during the division process. Of note, in some species, MreB relocalizes to the division site at the onset of cell constriction, suggesting that the elongasome and divisome cooperate during certain stages of the division cycle (Fenton and Gerdes, 2013; Figge et al., 2004).

While the function of the elongasome and divisome and their roles in establishment of generic rod and coccoid morphologies have been studied intensively (Egan et al., 2017; Typas et al., 2012), the mechanisms generating more complex cell shapes are still poorly understood. A model organism known for its distinctive morphological features is the alphaproteobacterium *Caulobacter crescentus* (henceforth *Caulobacter*) (Poindexter, 1964). This species is characterized by a biphasic life cycle that involves two morphologically and physiologically distinct cell types. One of them, the swarmer cell, possesses a single polar flagellum mediating swimming motility. The stalked cell, by contrast, displays a tubular extension (stalk) whose tip carries an adhesive holdfast mediating surface attachment. Whereas the stalked cell undergoes repeated cycles of chromosome replication and cell division, the swarmer cell is arrested in G1 phase, searching its environment for nutrients. However, at a defined point in the cell cycle, it sheds its flagellum, starts to establish a stalk at the previously flagellated pole, and enters S phase. The cell then elongates, forms a new flagellum at the pole opposite the stalk, and finally divides asymmetrically to produce a stalked cell and a new swarmer cell (Curtis and Brun, 2010). The biological role of the *Caulobacter* stalk is still controversial, but it may serve as a spacer to elevate the cell above the substratum and thus enhance its access to nutrients (Klein et al., 2013). Consistent with this idea, its length increases up to 20-fold under conditions of phosphate limitation (Schmidt and Stanier, 1966).

In *Caulobacter* species, the stalk consists almost exclusively of the three cell envelope layers (inner membrane, cell wall and outer membrane) and does not contain any cytoplasm (Ireland et al., 2002; Poindexter, 1964). Moreover, it is compartmentalized by large disc-like protein complexes, so-called crossbands, which are deposited at irregular intervals along its length, serving as non-selective diffusion barriers that physiologically separate the stalk envelope from the cell body (Poindexter, 1964; Schlimpert et al., 2012). Formation of the stalk is driven by zonal incorporation of new cell wall material at the stalk base, as detected by the labeling of newly synthesized PG with tritiated glucose (Schmidt and Stanier, 1966), radiolabeled D-cysteine (Aaron et al., 2007), or fluorescently labeled D-alanine derivatives (Kuru et al., 2012). To date, various mutants have been identified that lack stalks under standard growth conditions (Biondi et al., 2006; Bowman et al., 2008; Ebersbach et al., 2008; Sommer and Newton, 1989). However, in all cases, cells regained the ability to form stalks after transfer into phosphate-limited media, indicating that they suffered from a block in the cell cycle-regulated initation of stalk formation rather than a defect in the underlying biosynthetic machinery. By contrast, depletion of MreB or the hypothetical elongasome-specific GTase RodA (Meeske et al., 2016) was shown to impair stalk elongation under all growth conditions (Wagner et al., 2005). Similar results were obtained upon inhibition of the elongasome-specific TPase PBP2 (Seitz and Brun, 1998) with the β-lactam antibiotic mecillinam. However, because of the global effects of these treatments, it was difficult to conclude on a specific role of the three proteins in the stalk biosynthetic pathway. Finally, a moderate reduction in stalk length was observed for mutants lacking the cytoskeletal protein bactofilin A (BacA) or the BacA-associated class A PBP PbpC (Kühn et al., 2010). Together, these results suggest that components of the generic PG biosynthetic apparatus may be critical for stalk formation, but the precise composition of the machinery responsible for this process still remains elusive.

In the present study, we comprehensively investigate the mechanism of stalk formation, focusing on phosphate-limiting conditions to obtain a sensitive readout of the contributions that individual factors make to this process. We show that phosphate starvation induces a G0-like resting state that is characterized by the absence of key cell cycle regulators, including FtsZ. Comparing the muropeptide profiles of isolated stalk and cell body PG, we then identify significant differences in the composition of cell walls from these two compartments, suggesting that stalks are formed by specialized machinery with distinct biosynthetic properties. Systematic deletion and localization studies of cytoskeletal and PG biosynthetic proteins then indeed reveal a distinct set of factors involved in stalk elongation, which we characterize in detail with respect to their impact on PG composition and the spatial regulation of PG biosynthesis. Morphometric analysis of the corresponding mutants shows that these factors make varying and, in part specific, contributions to stalk and cell body elongation, indicating that these two modes of growth a mechanistically distinct. Finally, we identify MreB as a key component of the stalk biosynthetic complex and pinpoint a region on its surface that appears to be required for stalk formation but largely dispensable for elongasome-mediated lateral growth. Collectively, our results show that stalk formation represents a specialized growth process that is mediated by a composite complex including components of both the elongasome and divisome, with distinctive properties that clearly differentiate it from other PG biosynthetic machineries.

## Results

### Phosphate limitation arrests the cell cycle of *Caulobacter* in G1-phase

Although the stimulatory effect of phosphate starvation on *Caulobacter* stalk elongation has been known for decades (Schmidt and Stanier, 1966), the underlying regulatory mechanisms are still poorly understood. Prompted by the fact that stalk formation is tightly linked to cell cycle progression, we set out to investigate the effects of phosphate deprivation on central cellular processes such as DNA replication and cell division. First, flow cytometry was used to assess the replicational state of cells after transfer from standard to phosphate-free (M2G^-P^) medium. To this end, replication initiation was blocked with rifampicin and ongoing rounds of replication were allowed to finish. Previous work has shown that *Caulobacter* cells contain a single chromosome that is replicated only once per division cycle (Collier, 2012; Quon et al., 1998). Consistent with this finding, we observed that cells accumulated either one or two chromosome equivalents when grown in standard conditions, indicating that a large fraction of the population was in S-phase (**Figure 1A**). However, upon phosphate deprivation, DNA replication gradually ceased, with most cells arrested in G1-phase after 24 h of incubation. These data suggest that the lack of phosphate leads to a block in the cell cycle prior to S-phase, thereby preventing new rounds of chromosome replication. To support this conclusion, we visualized the number and positions of the chromosomal replication origins. In doing so, we made use of a fluorescently (GFP-) tagged derivative of the chromosome partitioning protein ParB, which interacts with specific motifs (*parS*) in the origin region (Mohl and Gober, 1997; Thanbichler and Shapiro, 2006). The expression of GFP-ParB thus typically results in the detection of either one or two foci, depending on the number of origin copies in the cell. Microscopic analysis revealed that most (^~^ 85%) cells exhibited a single ParB focus at the stalked pole when subjected to 24 h of phosphate starvation, indicating that they are arrested in G1 phase (**Figure 1B**). To clarify the reason for this G1 arrest, we analyzed the cellular levels of the replication initiator protein DnaA and the cell cycle master regulator CtrA, which act as positive and negative regulators of chromosome replication, respectively (Collier, 2012). Interestingly, both proteins were rapidly depleted from the cells during phosphate starvation (**Figure 1C**), indicating that key drivers of the *Caulobacter* cell cycle are absent under this condition.

**Figure 1.**
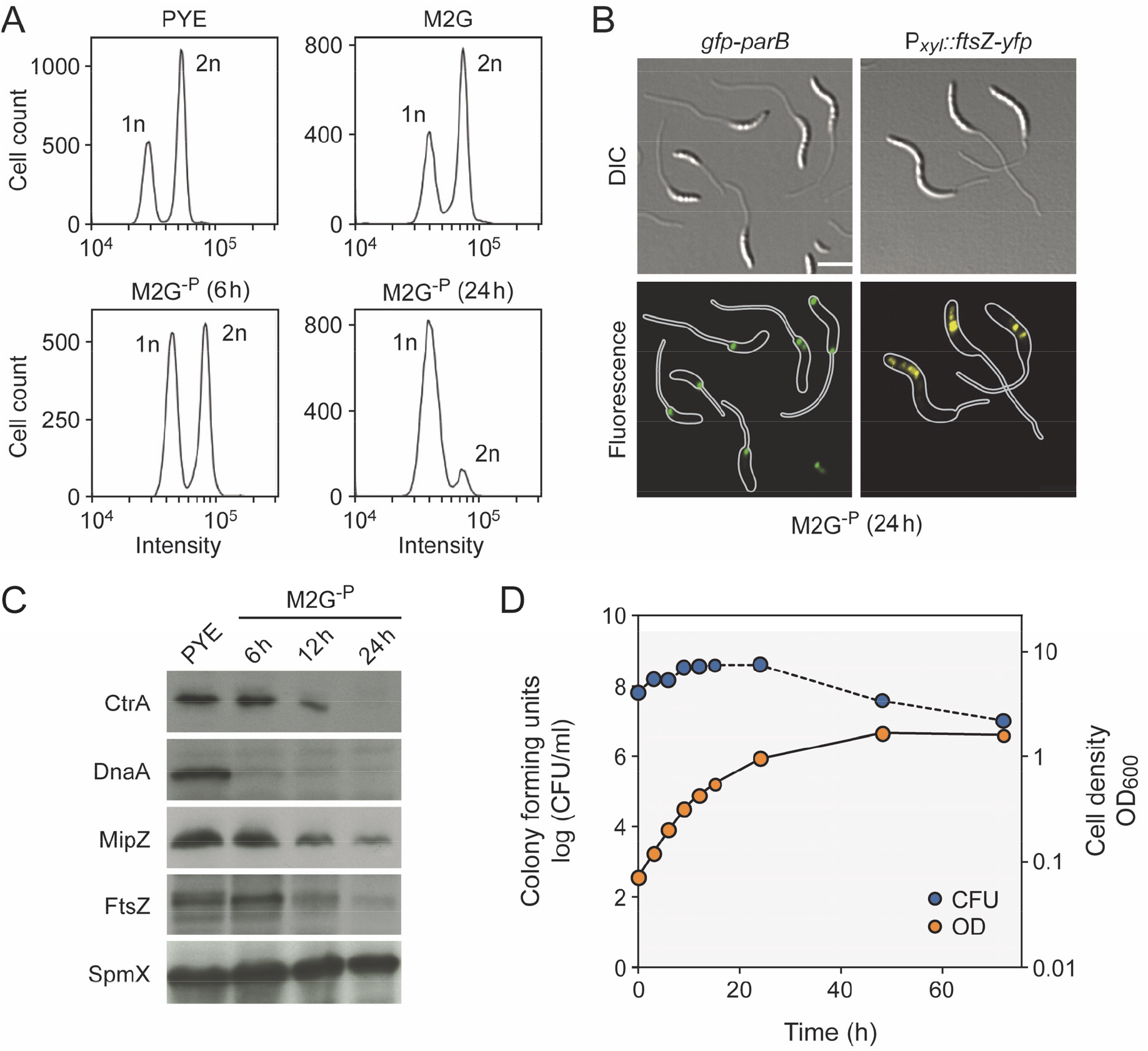
Progressive arrest of DNA replication and cell division under phosphate starvation. (**A**) DNA content of *C. crescentus* wild-type cells grown in PYE (rich medium, exponential phase), M2G (minimal medium, exponential phase), and M2G^-P^ (phosphate-lacking medium) for 12 h and 24 h. Cells were treated with 20 μg ml^-1^ rifampicin to prevent the reinitiation of replication prior to analysis by flow cytometry. (**B**) Subcellular localization of GFP-ParB and FtsZ-YFP in cells of strains MT199 (Pvan::Pvan-*ftsZ-yfp*) and MT174 (*parB*::*gfp-parB*) after 24 h of cultivation in M2G^-P^ medium. Synthesis of FtsZ-YFP was induced by addition of 50 μM vanillate 3 h prior to analysis (scale bar: 3 μm). (**C**) Changes in the levels of CtrA, DnaA, FtsZ, and MipZ over the course of phosphate starvation. Wild-type cells were grown in PYE, transferred into M2G^-P^ medium and subjected to Western blot analysis after 6 h, 12 h, and 24 h of incubation. A Western blot detecting SpmX served as a loading control. (**D**) Changes in the optical density (OD_600_) and viable-cell counts (CFU/ml) after transfer of a wild-type culture to M2G^-P^ medium.

To correlate changes in cell cycle progression with the growth behavior of cells, we monitored changes in cell mass and number after a shift to phosphate-limiting conditions. Interestingly, the optical density of cultures kept increasing exponentially for more than 10 h and only leveled off after ^~^ 50 h of incubation (**Figure 1D**), suggesting that cells made use of internal phosphate storage compounds to compensate for the lack of an external phosphate source. Consistent with the detection of DNA replication events (**Figure 1A**), cells still multiplied during the initial exponential phase. However, after longer starvation periods (> 24 h), the viable-cell count started to decline, whereas the cell mass still increased, likely due to continued elongation of the cell bodies and stalks in the absence of cell division events. Western blot analysis indeed revealed that the essential cell division protein FtsZ was depleted from the cells upon phosphate starvation (**Figure 1C**). The same was true for the cell division regulator MipZ, an inhibitor of FtsZ polymerization that limits Z-ring formation to the midcell region (Thanbichler and Shapiro, 2006). In line with these findings, an FtsZ-YFP fusion induced after prolonged phosphate starvation formed multiple foci in the vicinity of the stalk-distal pole instead of a defined midcell band (**Figure 1B**), indicating the absence of a functional and properly localized Z-ring (Thanbichler and Shapiro, 2006). Notably, FtsZ was never observed at the stalk base, supporting the previous notion that it does not play any role in stalk formation (Thanbichler and Shapiro, 2006).

Taken together, our results demonstrate that phosphate starvation arrests the *Caulobacter* cell cycle in a G1-like phase, thereby stalling DNA replication and cell division until phosphate becomes available again.

### Phosphate starvation induces a distinct pattern of PG synthesis

Phosphate starvation induces *Caulobacter* to enter a non-replicative resting state in which cells continue to elongate their cell body and stalk. To investigate this atypical mode of growth, we set out to visualize sites of active PG biosynthesis using the fluorescent D-amino acid 7-hydroxy-coumarin-amino-D-alanine (HADA) (Kuru et al., 2012; Kuru et al., 2015) as a tracer. As a control, we initially analyzed the growth dynamics of cells growing in phosphate-replete medium. To this end, cells were synchronized and then pulse-labeled with HADA at different stages of the cell cycle. Consistent with previous results (Aaron et al., 2007), we observed disperse incorporation of new cell wall material before the onset of cell division, followed by zonal growth at midcell during the constriction phase (**Figure 2–figure supplement 1**). Moreover, concurrent with the switch from disperse to zonal growth, an additional intense focus of fluorescence appeared at one of the cell poles, reflecting the establishment and outgrowth of the stalk. This polar signal faded gradually as the cell cycle progressed and was no longer detectable in late pre-divisional cells. Thus, HADA reliably detected all known growth zones in *Caulobacter* cells.

Next, we used HADA labeling to determine the pattern of PG synthesis under phosphate-limiting conditions (**Figure 2A**). After 6 h of incubation in phosphate-free medium, most cells showed a bright fluorescent patch at the stalked pole as well as a faint disperse signal extending throughout the rest of the cell body. Cells longer than ^~^ 4 μm often displayed an additional bright focus at their center, which likely reflects FtsZ-dependent zonal growth or cell division, consistent with the observation that the viable-cell counts still increased in the early phase of starvation (**Figures 1D** and **Figure 2–figure supplement 2A**). Interestingly, the intensity of the polar signal decreased considerably upon appearance of a midcell focus, suggesting that the machineries mediating stalk formation and cell division may compete with each other for at least some of their components (**Figure 2A**). After longer starvation periods (>18 h), midcell foci were almost undetectable, and HADA fluorescence was largely limited to the stalk base, which correlates with the lack of cell division events at this time point. Notably, the intensity of the polar signal decreased slightly during long-term incubation (**Figure 2–figure supplement 2B**), although the rate of stalk elongation remained constant at all time points (**Figure 2B**). The increase in cell body length, by contrast, was most pronounced during the early phases of starvation, when cells still showed midcell HADA foci, suggesting that it may, at least in part result, from FtsZ-mediated zonal growth at the cell center. Collectively, phosphate starvation induces a switch in the pattern of PG synthesis that ultimately limits cell growth to the stalked cell pole.

**Figure 2.**
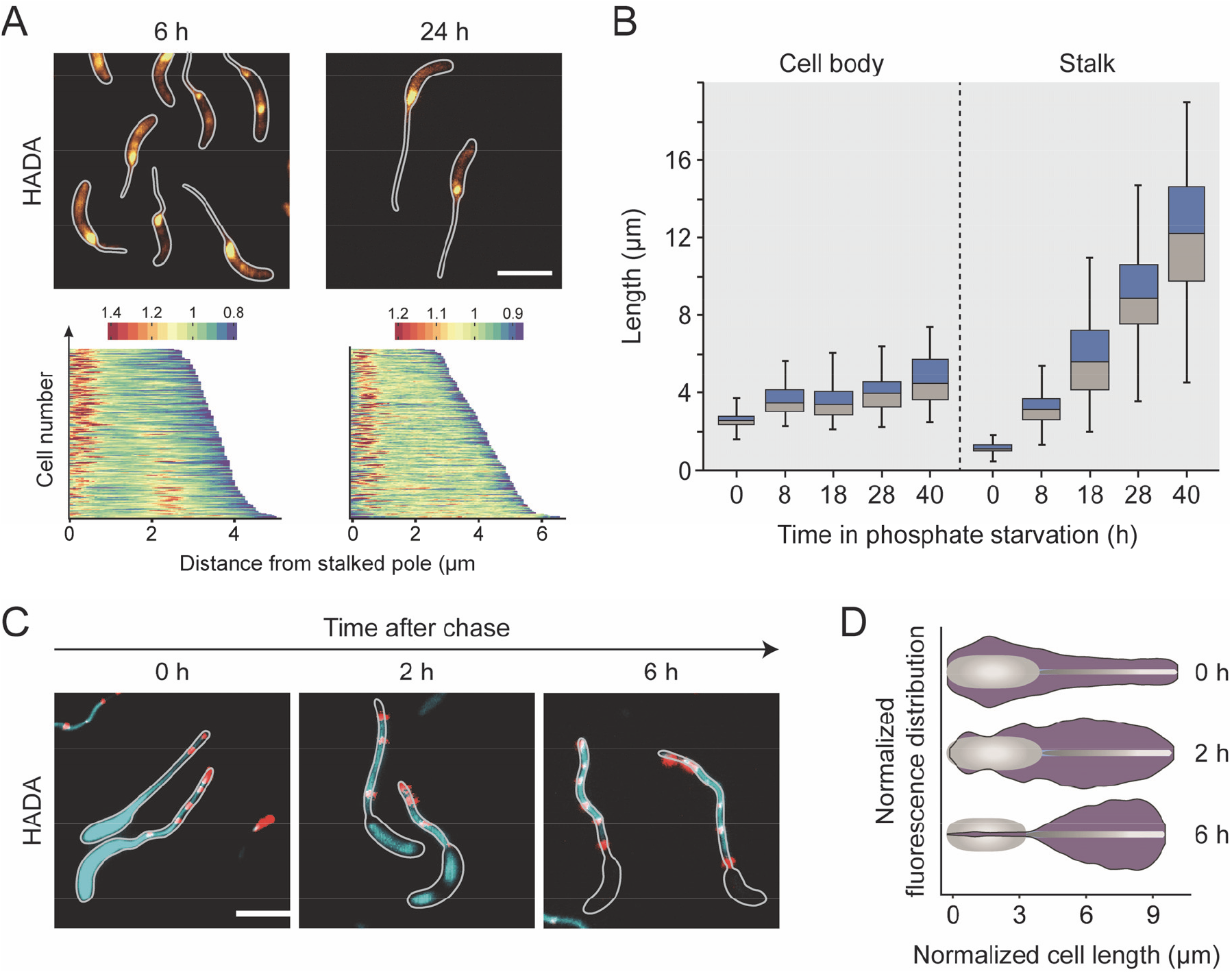
Reorganization of cell wall biosynthesis in the absence of phosphate. (**A**) Major growth zones of phosphate-starved wild-type cells. Cells were cultivated in M2G^-P^ medium for 6 h or 24 h and exposed to a short (2 min) pulse of HADA. The subcellular distribution of the fluorescence signals was quantified by demographic analysis of a random subpopulation of cells (n=200). To generate the graphs, single-cell fluorescence profiles were sorted according to cell length and stacked on top of each other (scale bars: 3 μm). (**B**) Changes in stalk and cell body lengths during phosphate starvation. Wild-type cells were incubated in M2G^-P^ medium for 8 h, 18 h, 28 h, and 40 h prior to imaging. The data are shown as box plots, with the horizontal line indicating the median, the box the interquartile range, and the whiskers the 2^th^ and 98^th^ percentile (0 h: n=210, 8 h: n=209, 28 h: n=208) (*** p < 10^-6^; t-test). (**C and D**) Slow turnover of PG in the stalk compartment. Cells were cultivated in M2G^-P^ medium for 18 h and exposed to HADA for an extended period of time (1.5 h). Subsequently, they were washed, transferred into fresh in M2G^-P^ medium and grown for 2 h, 4 h, and 6 h in the absence of the label (scale bars: 3 μm). To quantify the changes in HADA fluorescence overtime, fluorescence profiles were obtained from random subpopulations of cells (n=200 per time point). The lengths of the profiles in each quintile of the cell length distribution were normalized to the maximum cell length in the respective quintile. Subsequently, the fluorescence intensities were averaged and used to generate violin plots. Shown is a representative part of the data depicting the fluorescence distributions in the fourth quintile at each of the time points (**D**). The full analysis is presented in **Figure S2C**.

Our and previous labeling studies suggest that stalk formation is driven by the insertion of new cell wall material at the stalk base (Aaron et al., 2007; Kuru et al., 2012; Schmidt and Stanier, 1966). To determine whether stalk PG is still subject to modification or turnover, phosphate-starved *Caulobacter* cells were incubated with HADA for an extended period of time (1.5 h). After this treatment, staining was observed throughout the entire cell envelope, including distal segments of the stalk that were clearly formed prior to the start of the labeling procedure (**Figure 2C**). This finding demonstrates the presence of transpeptidase activity in the stalk compartment that mediated the incorporation of HADA independently of the pole-associated biosynthetic complex. After transfer of the cells to HADA-free medium, fluorescence was rapidly lost in the cell bodies and in the basal region of the stalk, whereas it was stably retained in the distal stalk segments, indicating that stalk PG is not turned over at significant rates (**Figure 2C**). Notably, the same behavior was observed for a strain lacking crossbands. The distinct behavior of cell bodies and stalks may thus not be due to restrictions in the diffusion of envelope-localized PG biosynthetic enzymes but, potentially, rather due to the absence of cytoplasm in the stalk compartment.

### The stalk and cell body show different PG architectures

The distinct mode of growth involved in stalk formation opens the possibility that there may be compositional differences between the PG layers encompassing the cell body and stalk compartments. To address this issue, phosphate-starved cells were agitated vigorously to shear off stalks from the cell bodies. After separation of the two compartments by differential centrifugation (**Figure 3–figure supplement 1**), PG was isolated from each of the fractions and subjected to muropeptide analysis. Interestingly, stalk PG contained a high proportion of 3-3 crosslinked peptides and non-crosslinked tripeptides (resulting from the cleavage of 3-3 bonds), whereas these muropeptide species were barely detectable in the cell body samples (**Figure 3** and **Supplementary file 1**). Similarly, the total fraction of crosslinked peptide side chains was significantly higher in stalk PG, mostly because of a higher proportion of trimeric muropepides. The glycan chain lengths, by contrast, did not vary between the two compartments. Collectively, these findings indicate that the PG layers of stalks and cell bodies differ in both the type and extent of peptide crosslinks.

**Figure 3.**
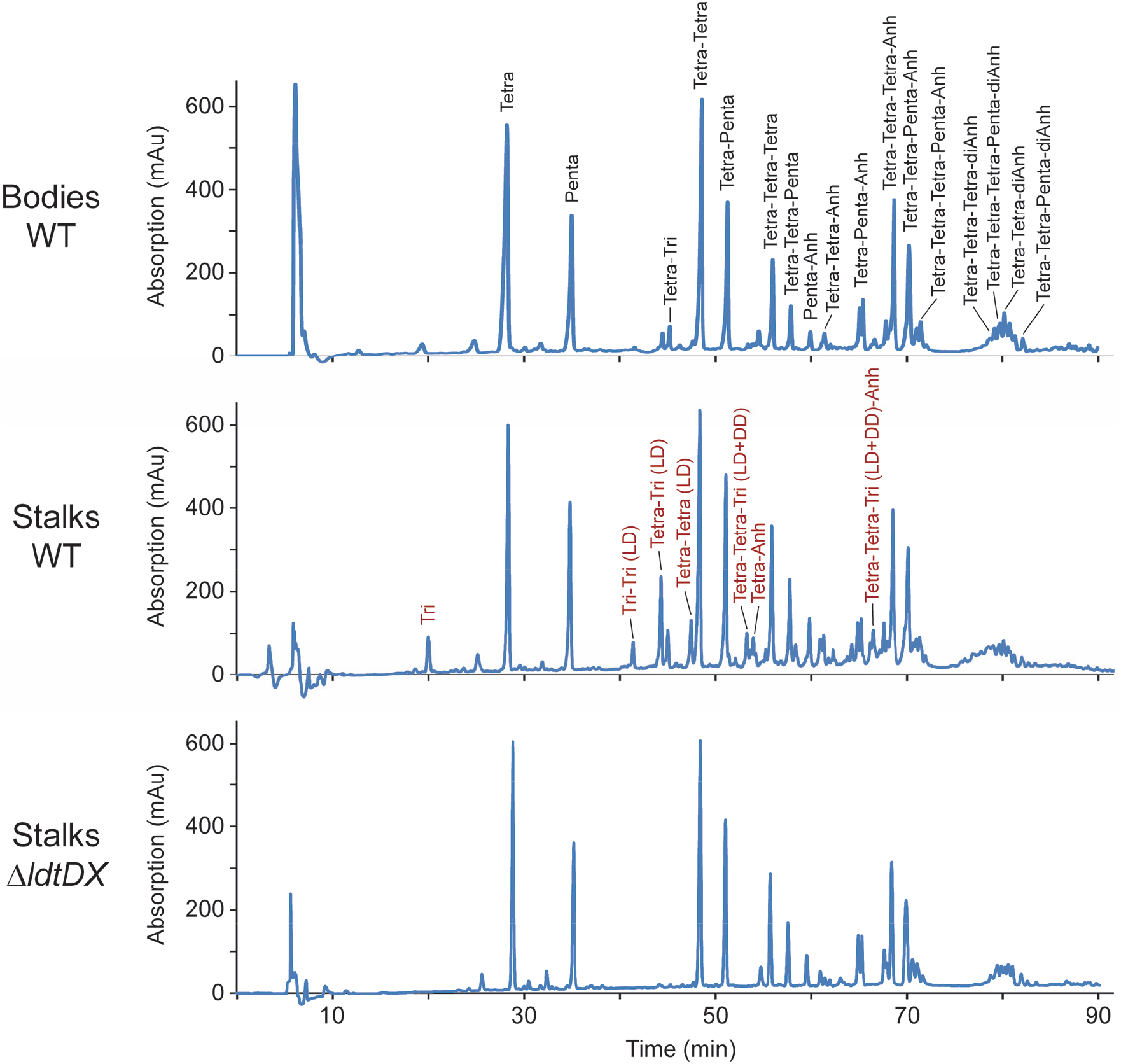
Differential composition of cell body and stalk peptidoglycan. Shown are the HPLC profiles of muropeptides obtained from the cell body and stalk fractions of stains NA1000 (WT) and AZ138 (Δ*ldtD* Δ*ldtX*) after growth in M2G^-P^ medium for 24 h. In the first panel, the identities of the most abundant muropeptides are given in black. In the second panel, products that are specifically enriched in the stalk fraction are indicated in red. Abbreviations: Tri: Glc*N*Ac–Mur*N*Ac(r)–L-Ala–D-Glu–mDap; Tetra: Glc*N*Ac–Mur*N*Ac(r)–L-Ala–D-Glu–mDap-D-Ala; Penta: Glc*N*Ac–Mur*N*Ac(r)–L-Ala–D-Glu–mDap-D-Ala–D-Ala; Anh: 1,6-anhydro-Mur*N*Ac; DD: mDap–D-Ala cross-link; LD: mDap–mDap crosslink.

Previous work has shown that 3-3 crosslinks are generated by LD-TPases, which are characterized by a conserved YkuD domain (Magnet et al., 2008). The *Caulobacter* genome contains two so-far uncharacterized open reading frames, CC_1511 and CC_3744, which encode proteins with this signature domain (now referred to as LdtD and LdtX, respectively). To determine how these factors contribute to the distinctive composition of the stalk cell wall, we generated a strain carrying in-frame deletions in both the *ldtD* and *ldtX* gene and analyzed the composition of PG purified from its stalk and cell body compartments. In both samples, 3-3 crosslinked peptides and non-crosslinked tripeptides were virtually undetectable (**Figure 3** and **Supplementary file 1**), indicating that the formation of these muropeptide species is linked to the activity of the two predicted LD-TPases. Notably, however, the total fraction of crosslinked peptides barely changed in either of the compartments, because the loss of 3-3 crosslinks was compensated by a proportional increase in the fraction of 4-3 crosslinks. Thus, LD-TPase activity is not the main factor responsible for the elevated degree of crosslinking in stalk PG.

Previous work has shown that the stalk is physiologically separated from the cell body, because it is devoid of cytoplasm and contains crossband complexes that block the exchange of periplasmic and membrane proteins (Schlimpert et al., 2012). It was conceivable that crossbands could help establish the differences in the PG composition observed for the two compartments, for instance by facilitating the establishment of distinct pools of PG biosynthetic enzymes or blocking the diffusion of lipid II into the stalk structure. To test this idea we determined the muropeptide profile of stalk and cell body PG isolated from a crossbandless strain (*ΔstpAB*, SW51). Notably, we still observed a higher content of 3-3 crosslinks and a higher total proportion of crosslinked peptides in stalk PG (**Supplementary file 1**). Similar to the differences in PG turnover (**Figure 2C**), this characteristic thus appear to be independent of the presence of crossbands.

### Stalk formation involves class A and class B penicillin-binding proteins

Stalk formation involves a growth process that is distinct from the disperse and zonal incorporation of PG mediated by the elongasome or division complex, respectively. To determine the composition of the underlying machinery, we systematically analyzed all predicted PG biosynthetic proteins encoded in the *Caulobacter* genome for their contribution to stalk elongation under phosphate-limiting conditions. In doing so, we initially focused on enzymes with PG synthase activity, including PBPs and LD-TPases. A previous study has shown that inhibition of the monofunctional DD-TPase PBP2 with mecillinam largely abolished the synthesis of stalks under standard conditions, although it concomitantly induced severe morphological defects in the cell body (Seitz and Brun, 1998). To further investigate the role of this protein, we generated a strain producing a fully functional GFP-PBP2 fusion in place of the wild-type protein and analyzed the localization pattern of the fluorescently tagged protein under conditions of phosphate starvation. The majority of cells, in particular those with clearly elongated (longer than ^~^ 4 μm) cell bodies, showed a faint focus at the stalked cell pole (**Figure 4A**), supporting the idea that PBP2 may have a role in stalk formation. Previous work has also implicated bifunctional PBPs in stalk elongation, largely based on the analysis of mutants lacking one of these proteins (Kühn et al., 2010; Yakhnina & Gitai, 2013). To verify and extend these results, we analyzed strains carrying single or multiple mutations in the PBP-encoding *pbpY, pbp1A, pbpC, pbpX*, and *pbpZ* genes (Strobel et al., 2014; Yakhnina and Gitai, 2013). Our results confirm that deletion of *pbpC* led to a moderate reduction in stalk length, whereas the absence of any other PBP, either alone or in combination, did not have any effect (**Figure 4B**). However, as observed under standard growth conditions (Strobel et al., 2014; Yakhnina and Gitai, 2013), at least one bifunctional PBP was required for viability during phosphate starvation (**Figure 4–figure supplement 1A**). In line with the results of the deletion studies, localization analyses revealed that none of the bifunctional PBPs except for PbpC accumulated at the stalked pole, indicating that these proteins may not be specifically associated with the stalk biosynthetic machinery (**Figure 4–figure supplement 1B**). Notably, however, PbpX appeared enriched in the stalk compartments, but the significance of this observation remains unclear.

**Figure 4.**
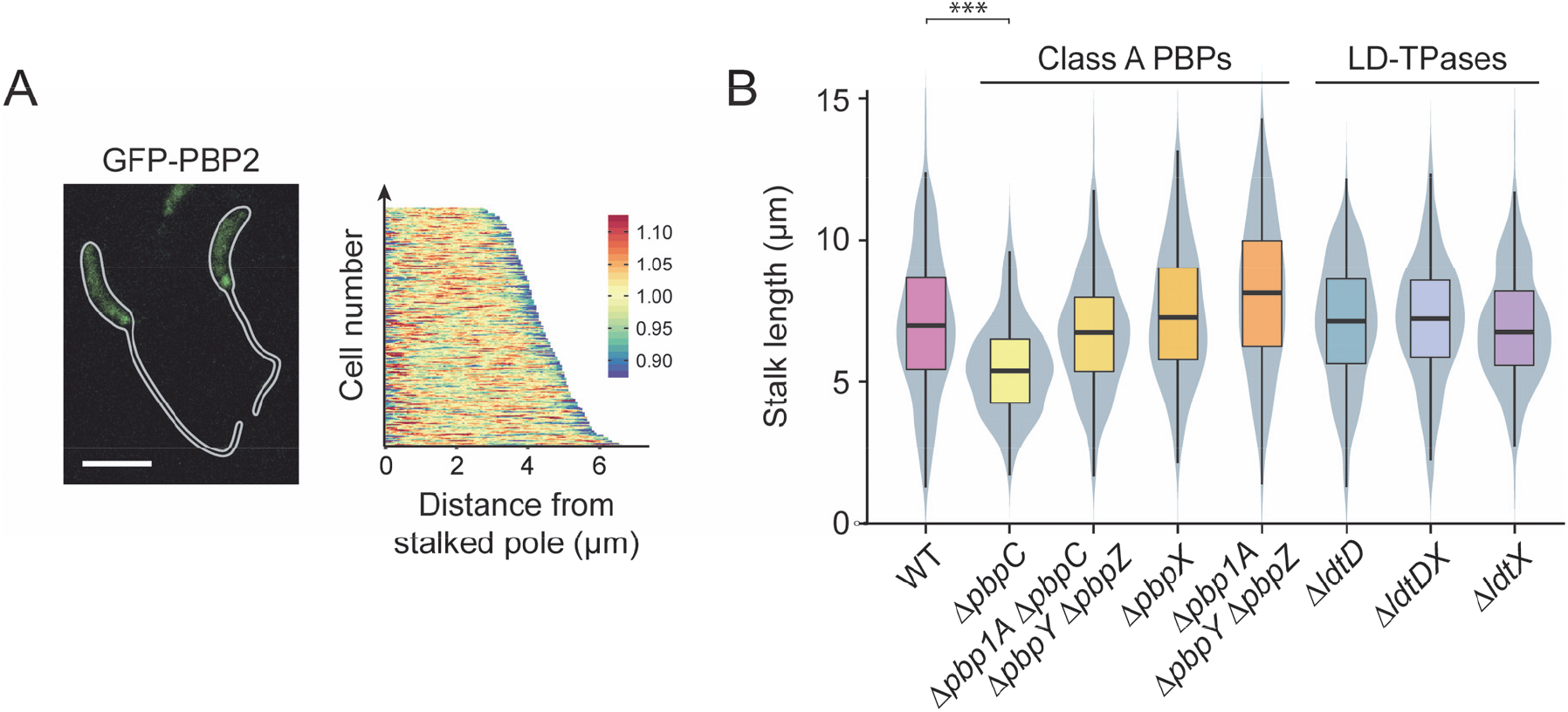
Participation of PG biosynthetic enzymes in stalk elongation. (**A**) Localization of GFP-PBP2 fusion in strain MAB244 (*pbp2::gfp-pbp2*) after 24 h of cultivation in M2G^-P^. The demograph shows the fluorescence profiles of a random subpopulation of cells sorted according to cell length (n=227). (**B**) Distribution of the stalk lengths in populations of mutants lacking specific PG synthases. Shown are the results obtained for MT286 (Δ*pbpC*), JK305 (Δ*pbpA1* Δ*pbpC* Δ*pbpY* Δ*pbpZ*), KK1 (Δ*pbpX*), KK12 (Δ*pbp1A* Δ*pbpY* Δ*pbpZ*), AZ137 (Δ*ldtD*), AZ138 (Δ*ldtDX*), and AZ140 (Δ*ldtX*) after 24 h of growth in M2G^-P^ medium. Data are represented as box plots, with the horizontal line indicating the median, the box the interquartile range and the wiskers the 2^nd^ and the 98^th^ percentile (n=210 per strain). In addition rotated kernel density plots (grey) are depicted for each dataset to indicate the distribution of the raw data (*** p < 10^-6^; t-test).

Finally, we analyzed the role of the two predicted LD-TPases LdtD and LdtX in stalk formation. Although these proteins make a significant contribution to PG crosslinking in the stalk compartment (**Figure 3**), their inactivation did not have any apparent phenotypic effect (**Figure 4B**). LD-TPase activity may thus not contribute to the establishment of the stalk structure per se but rather have an accessory function that serves to modify the biophysical properties of the PG layer. Localization studies indicate that LdtD and LdtX do not accumulate at the stalk base, suggesting that they may act independently of the polar stalk biosynthetic machinery (**Figure 4–figure supplement 1C**).

### Components of the autolytic machinery are critical for stalk formation

Apart from PG synthases, stalk formation must also involve autolytic enzymes that cleave the PG sacculus and, thus, enable the insertion of new cell wall material at the stalk base. However, to this point, the nature of the factors involved has remained unknown. To address this issue, we systematically screened mutants lacking one or multiple predicted PG hydrolases for defects in stalk growth under phosphate-limiting conditions. The enzymes tested included all LytM-like and NlpC/P60-like endopeptidases, AmiC-like and CHAP domain-containing amidases, soluble and membrane-bound lytic transglycosylases, and carboxypeptidases identified in the *Caulobacter* genome (**Supplementary file 2**). In most cases, the lack of single factors and even the absence of whole enzyme families had no apparent effect on stalk length (**Figure 5–figure supplement 1**). Four strains, however, displayed obvious morphological defects (**Figure 5**). One of them was a mutant lacking the protein DipM, a catalytically inactive LytM-like endopeptidase homolog that was previously shown to be critical for proper PG remodeling during cell division (Goley et al., 2010; Möll et al., 2010; Poggio et al., 2010). The absence of DipM led to a severe reduction in stalk length, combined with the formation of branches within the stalk structure or the establishment of multiple stalks, often emanating from the same pole (**Figure 5–figure supplement 2**). Even shorter stalks were observed in the combined absence of the soluble lytic transglycosylases SdpA and SdpB, two proteins previously found to be associated with the divisome complex (Zielinska et al., 2017). Apart from its aberrant morphology, the *ΔsdpAB* mutant frequently showed membrane blebs that were associated with the residual stalk structures, suggesting a defect in membrane attachment or homeostasis (**Figure 5–figure supplement 2**). Milder effects on stalk length were caused by inactivation of the divisome-associated carboxypeptidase CrbA (Billini *et al*, unpublished) (**Figure 5–figure supplement 2**) and the LytM-like endopeptidase LdpA, a thus-far uncharacterized protein encoded in an operon with the polarly localized scaffolding protein bactofilin A (BacA) (Kühn et al., 2010; Shi et al., 2015) (**Figure 5–figure supplement 2**). Importantly, despite their defects in stalk elongation, none of the four strains showed a significant reduction in cell length (**Figure 5C**), indicating that stalk and cell body growth are mechanistically distinct processes that proceed independently of each other.

**Figure 5.**
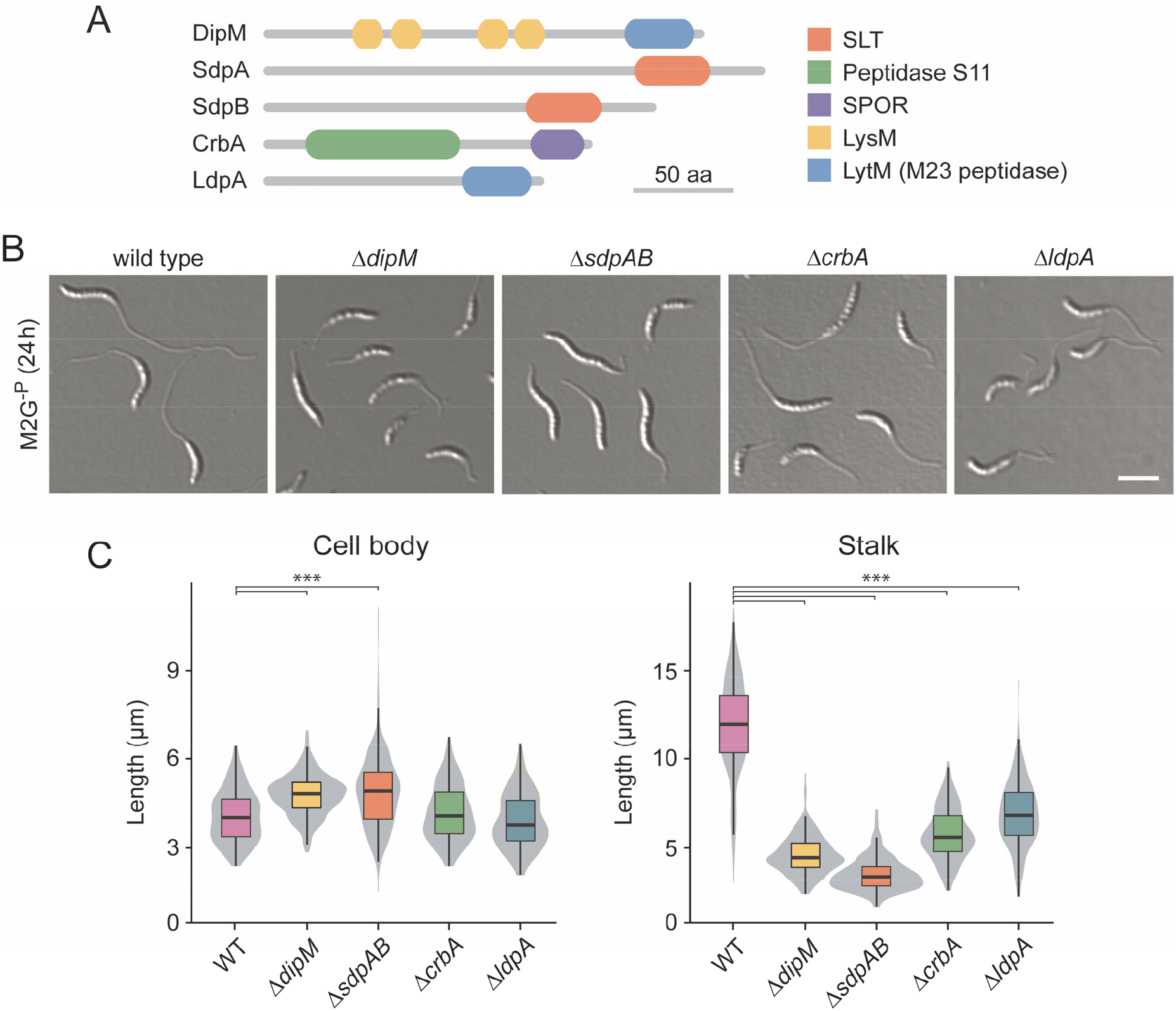
Participation of autolytic factors in stalk elongation. (**A**) Domain structure of selected components of the autolytic machinery of *C. crescentus*. (**B**) DIC micrographs of mutant cells exhibiting a stalk elongation defect. Shown are strains MT258 (Δ*dipM*), AZ22 (Δ*sdpAB*), AM376 (Δ*crbA*), and AM364 (Δ*ldpA*) in comparison to NA1000 (WT) after 24 h of cultivation in M2G^-P^ medium. (**C**) Distribution of the cell body and stalk lengths in populations of strains MT258, AZ22, AM376, and AM364 after growth in M2G^-P^ for 24 h. The values obtained are shown as box plots, with the horizontal line indicating the median, the box the interquartile range and the wiskers the 2^nd^ and the 98^th^ percentile (n=210 per strain). In addition rotated kernel density plots (grey) are depicted for each dataset to indicate the distribution of the raw data (*** p < 10^-6^; t-test).

To further investigate the functions of the five autolytic factors identified in the mutational screen, we generated fluorescently (mCherry-) tagged derivatives of these proteins and analyzed their localization patterns under conditions of phosphate starvation (**Figure 6**). Both the DipM and CrbA fusions accumulated at the stalk base and may, thus, be specifically associated with the polar stalk biosynthetic machinery. The SdpA and SdpB fusions, by contrast, were distributed throughout the cell envelope, suggesting that the two proteins may either act independently of the polar complex or associate with it in a very transient manner. Unlike the other proteins analyzed (**Figure 6–figure supplement 1**), LdpA-mCherry was quantitatively cleaved at the junction between the two fusion partners, preventing further analysis (data not shown).

**Figure 6.**
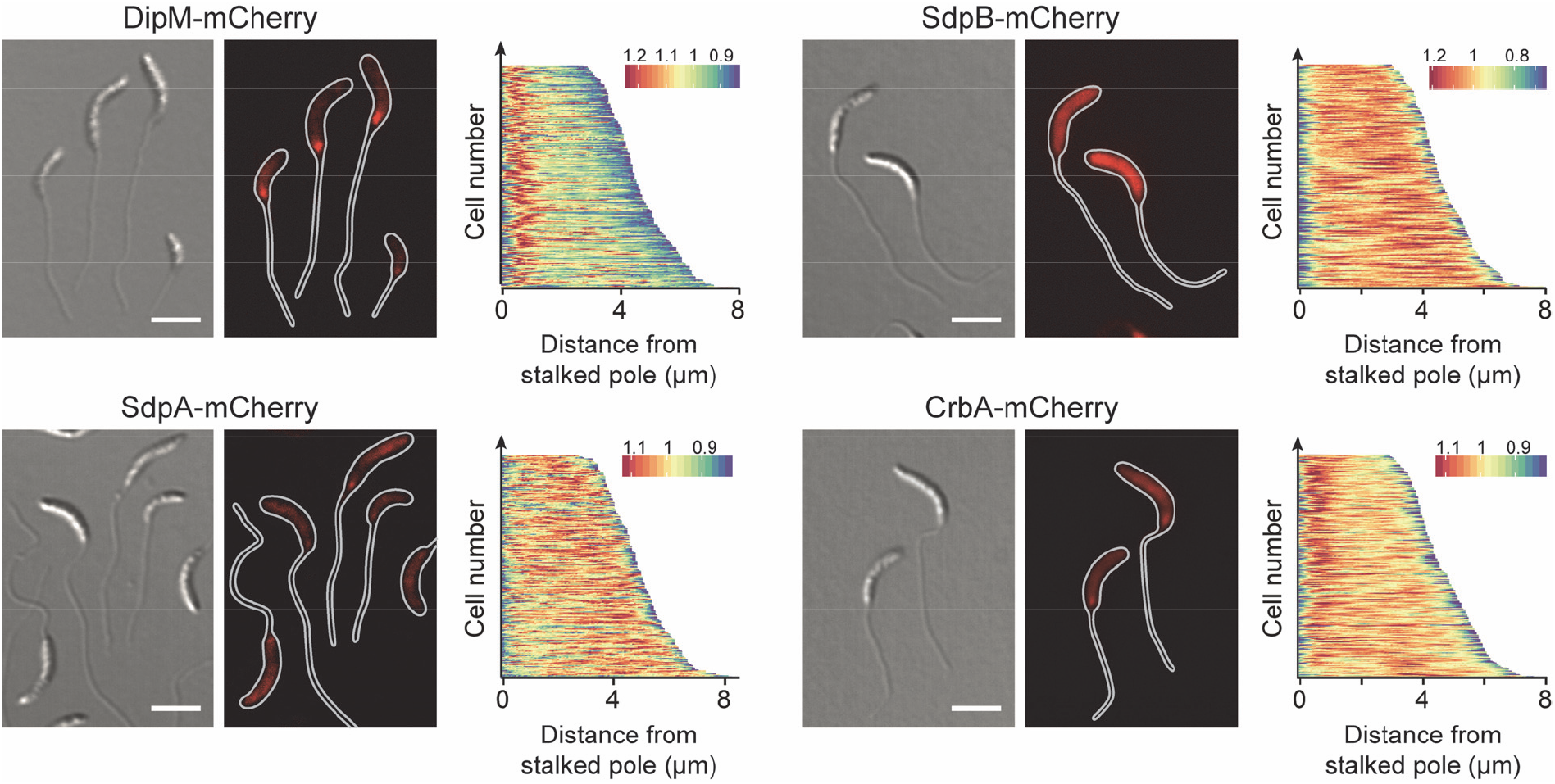
Localization of autolytic factors in phosphate-starved cells. Shown are the localization patterns of SdpA-mCherry (AM480, Pxyl::Pxyl-*sdpA-mCherry*), CrbA-mCherry (MAB247, Pxyl::Pxyl-*crbA-mCherry*), DipM-mCherry (AM208, Pxyl::Pxyl-*dipM-mCherry*), and SdpB-mCherry (AZ127, Pxyl::Pxyl-*sdpB-mCherry*) in cells cultivated for 24 h in M2G^-P^ medium (scale bars: 3 μm). Synthesis of the fluorescent protein fusions was induced for 3 h (for DipM, SdpA, and CrbA) or 2 h (for SdpB) with 0.3% xylose prior to analysis. The demographs next to the images show the fluorescence profiles of a random subpopulation of cells sorted according to cell length (n=200 for each strain).

In order to determine how the absence of the different autolytic factors influenced the pattern of PG biosynthesis, mutants lacking these proteins were grown in phosphate-limiting conditions and subjected to HADA staining (**Figure 7**). Consistent with their relatively mild stalk elongation defect, *ΔldpA* cells still displayed a pattern similar to that of the wild-type strain. In the *ΔdipM* and *ΔcrbA* strains, by contrast, the polar signals were much fainter and new cell wall material was often incorporated at non-polar sites. An even more pronounced effect was observed in the *ΔsdpAB* mutant, which virtually lacked polar foci and instead showed patchy or even HADA fluorescence throughout the cells. Thus, the severity of the stalk elongation defect scales with the loss in polar PG biosynthesis.

**Figure 7.**
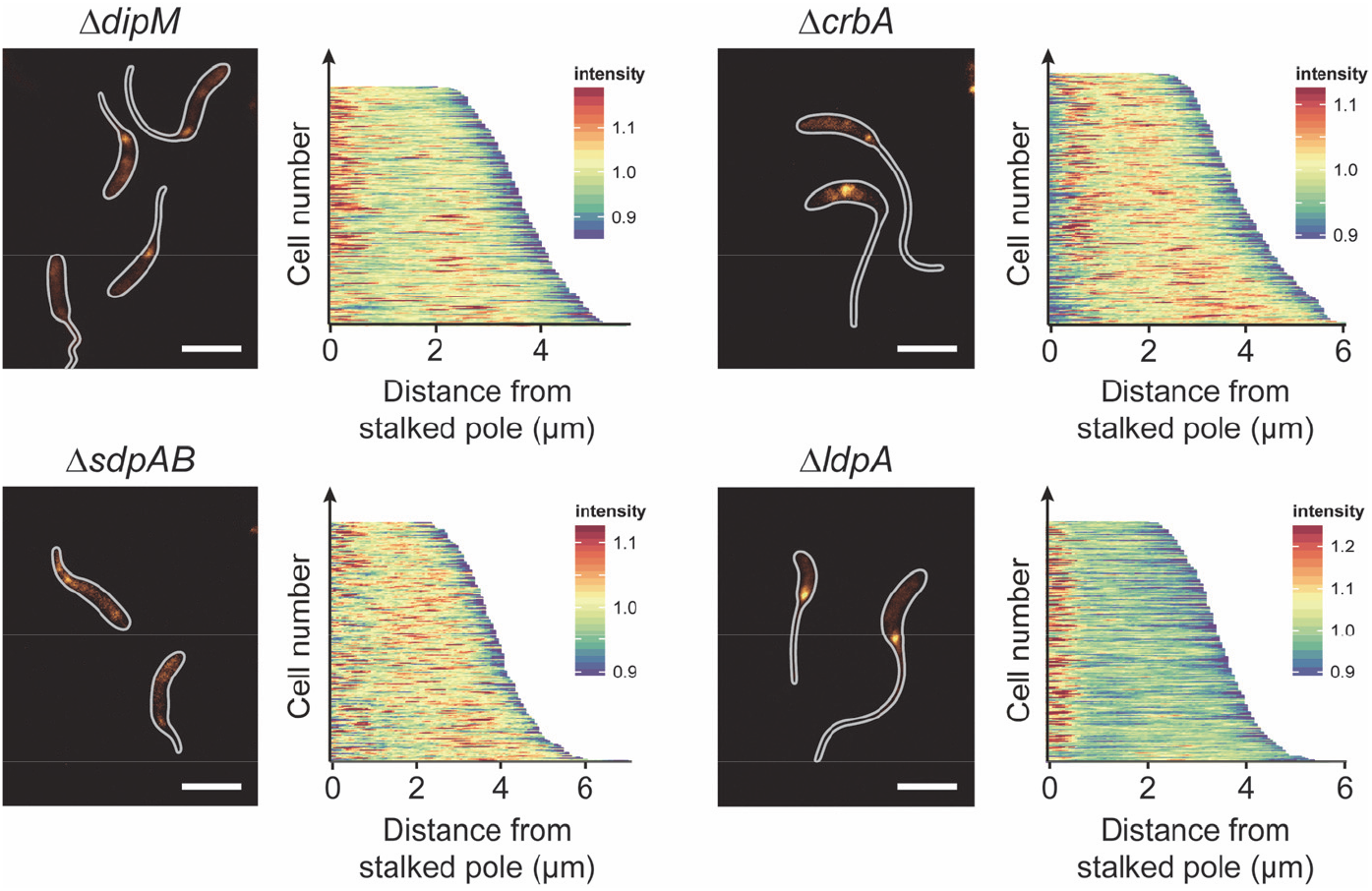
Cell wall biosynthesis in mutants with defects in the autolytic machinery. Shown are fluorescence images of strains NA1000 (WT), MT258 (Δ*dipM*), AZ22 (Δ*sdpAB*), AM376 (Δ*crbA*), and AM364 (Δ*ldpA*) after 24 h of incubation in M2G^-P^ medium and subsequent HADA staining (2 min). The distribution of the fluorescence signals was quantified by demographic analysis of a random subpopulation of cells (n=200 for each strain) (scale bars: 3 μm).

To obtain more detailed insight into the effects of the different mutations on the structure of the PG layer, we isolated whole-cell sacculi from wild-type and mutant cells after prolonged (24 h) phosphate starvation and subjected them to muropeptide analysis (**Supplementary file 3**). For the wild-type strain, wholecell sacculi gave similar results as PG from isolated from a cell body fraction (compare **Figure 3** and **Supplementary file 1**), indicating that the characteristic features of stalk PG are largely obscured by the excess of cell body PG in the whole-cell preparations. Interestingly, there were hardly any differences between the muropeptide profiles obtained under phosphate-limiting (**Supplementary file 3**) and phosphate-replete (Takacs et al., 2010) conditions. The average composition of cell body PG thus appears to be independent of the phosphate supply. Among the mutant strains, *ΔldpA* cells showed essentially the same average PG composition as the wild-type strain. The same was true for the *ΔdipM* mutant, with exception of a significant increase in the proportion of non-crosslinked tetrapeptides (**Supplementary file 3**), which could indicate an elevated level of endopeptidase and/or carboxypeptidase activity. The muropeptide profiles of the remaining strains, by contrast, showed marked global changes. In line with the notion that CrbA acts as a carboxypeptidase, removing the terminal D-Ala residue of pentapeptide side chains (Billini *et al*, unpublished), the *ΔcrbA* mutant displayed a considerable decrease in the total content of tetrapeptides that was accompanied by a proportional increase in the content of pentapeptide-containing muropeptide species (**Supplementary file 3**). In *ΔsdpAB* cells, on the other hand, the average glycan chain length increased from 7 to 9.4 disaccharide units, consistent with the loss of lytic transglycosylase activity. Surprisingly, the mutant cells additionally showed a severe reduction in the degree of crosslinkage. At the same time, their total content of pentapeptide side chains was reduced, whereas the proportion of tripeptide side chains was considerably elevated (**Supplementary file 3**). These results suggest that the lack of SdpAB leads to reduced transpeptidation or, more likely, elevated endopeptidase activity.

Collectively, our results show that several components of the autolytic machinery are critical for proper PG remodeling during stalk formation, with some of them localizing to the stalk base under phosphate-limiting conditions. Notably, most of the proteins, including DipM, SdpA, SdpB and CrbA, are associated with the cell division apparatus under standard growth conditions {Möll et al., 2010; Poggio et al., 2010; Goley et al., 2010; Zielinska et al., 2017; M. Billini, unpublished), suggesting parallels in the mechanisms underlying cell constriction and stalk growth.

### Stalk formation depends on the presence of scaffolding proteins

Polymer-forming scaffolding proteins are critical for the regulation of many growth processes in bacteria {den Blaauwen et al., 2008; Lin and Thanbichler, 2013}, suggesting that this group of proteins may also play a critical role in stalk formation. Previous work has indeed implicated the bactofilin homolog BacA in stalk biogenesis (Kühn et al., 2010). Re-analysis of a *ΔbacA* mutant revealed a significant reduction in both stalk and cell body length during phosphate starvation (**Figures 8** and **Figure 8–figure supplement 1**). Notably, deletion of the endopeptidase gene *ldpA*, which lies in a putative operon with *bacA*, had a very similar effect on stalk length, whereas it barely affected the cell body (**Figure 5**). These results suggest that LdpA and BacA may specifically cooperate in stalk formation, whereas BacA is additionally involved in a distinct pathway involved in cell body elongation.

**Figure 8.**
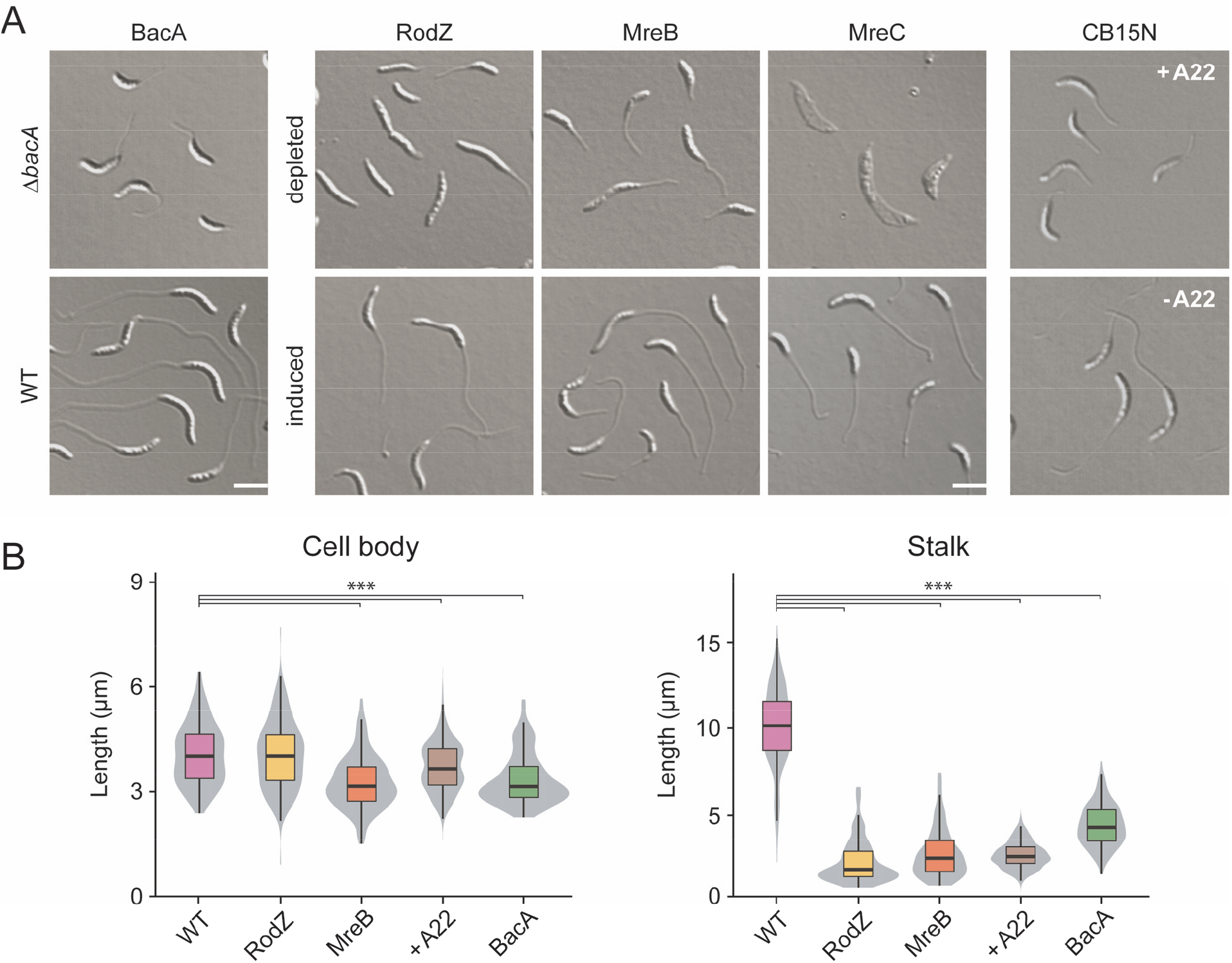
Role of scaffolding proteins in stalk elongation. (**A**) DIC images of cells lacking single scaffolding proteins. The strains analyzed were LS4275 (Δ*mreC* Pxyl::Pxyl-*mreC*), LS3809 (Δ*mreB* Pxyl::Pxyl-*mreB*), CJW2747 (Δ*rodZ::Ω* Pxyl::Pxyl-*rodZ*), MT257 (Δ*bacA*). Strain NA1000 (WT) is shown as a wild-type control. Strains LS4275, LS3809, and CJW2747 were grown to exponential phase PYE medium containing the inducer xylose. Subsequently, the cells were washed, grown for another 7 h in PYE medium with or without inducer and then incubated for 24 h in M2G^-P^ medium with or without inducer prior to imaging by DIC microscopy. Strain MT257 was grown to stationary phase in PYE medium, diluted (1:20) into M2G^-P^ medium, and grown for 24 h prior to analysis. Strain NA1000 (WT) was grown in M2G^-P^ medium. After 9 h, the cultures were supplemented with A22 at a final concentration of 10 μg/ml and incubated for additional 15 h prior to imaging (scale bar: 3 μm). (**B**) Distribution of cell body and stalk lengths in populations of WT NA1000 and depleted strains LS4275, LS3809, CJW2747, and MT257 grown as described in (**A**). The data are shown as box plots, with the horizontal line indicating the median, the box the interquartile range and the wiskers the 2^nd^ and the 98^th^ percentile (n=210 per strain). In addition rotated kernel density plots (grey) are depicted for each dataset to indicate the distribution of the raw data (*** p < 10^-6^; t-test).

As another scaffolding protein, MreB was shown to be required for stalk formation in media containing moderate to high levels of phosphate (Divakaruni et al., 2007; Wagner et al., 2005). To clarify the contribution of this protein to stalk biosynthesis under phosphate starvation, we employed strains producing MreB or the adapter protein RodZ under the control of an inducible promoter. When starved for phosphate in the absence of inducer, both mutants showed a drastic reduction in stalk length or occasionally even failed to form stalks at all (**Figures 8 and Figure 8– figure supplement 1**). The effects on the cell bodies, by contrast, differed depending on the protein depleted. Cells lacking RodZ showed a length distribution indistinguishable from that of the wild-type strain. Depletion of MreB, by contast, markedly decreased the median cell length. Similar effects were observed for wild-type cells treated with the MreB inhibitor A22 (Gitai et al., 2005; van den Ent et al., 2014) (**Figure 8**). These findings indicate that, under phosphate-limiting conditions, RodZ appears to be specifically required for stalk biosynthesis, whereas MreB additionally contributes to cell body elongation, again supporting the idea that these two processes are driven by distinct mechanisms.

Apart from MreB, the MreCD complex has been identified as a factor critical to lateral growth in many rod-shaped bacteria (den Blaauwen et al., 2008). MreC is thought to serve as a scaffold that interacts with various PG biosynthetic enzymes, including the monofunctional TPase PBP2 (Contreras-Martel et al., 2017; Divakaruni et al., 2005). In *E. coli*, it is part of the elongasome complex (Kruse, Bork-Jensen, & Gerdes, 2005), whereas it was shown to establish an elongasome-independent structure in *Caulobacter* cells (Divakaruni et al., 2007; Dye et al., 2005). To test for a role of this protein in stalk formation, we analyzed the morphology of a conditional *mreC* mutant grown under phosphate-limiting conditions (**Figure 8**). In the absence of inducer, the cells started to elongate but eventually became amorphous and lyzed. In most cases, stalks were either absent or barely recognizable, indicating that the MreCD complex may be essential for both cell wall integrity and stalk biosynthesis during phosphate starvation. Collectively, our results support the notion that various scaffolding proteins are required for proper stalk biosynthesis in *Caulobacter* cells.

To clarify whether the role of the different scaffolding proteins in stalk formation involves their recruitment to the stalked pole, we analyzed the localization patterns of fluorescently tagged derivatives in cells subjected to phosphate starvation (**Figure 9**). Both the MreB and RodZ fusion formed a distinct focus at the stalk base and, in rare cases, also a second focus at the pole opposite the stalk. Together with the polar localization of PBP2 (**Figure 4A**), these findings indicate that key components of the elongasome complex relocate to the site of stalk biosynthesis in phosphate-limiting conditions. There, they colocalize with BacA, which retains its polar position irrespective of changes in the phosphate supply (**Figure 9**). The MreC fusion, by contrast, formed a broad band at midcell, whereas it was largely excluded from the polar regions (**Figure 9**). In line with the global morphological defects caused by its depletion, MreC may have a general role in cell wall integrity, but it does not appear to be specifically associated with the polar stalk biosynthetic machinery.

**Figure 9.**
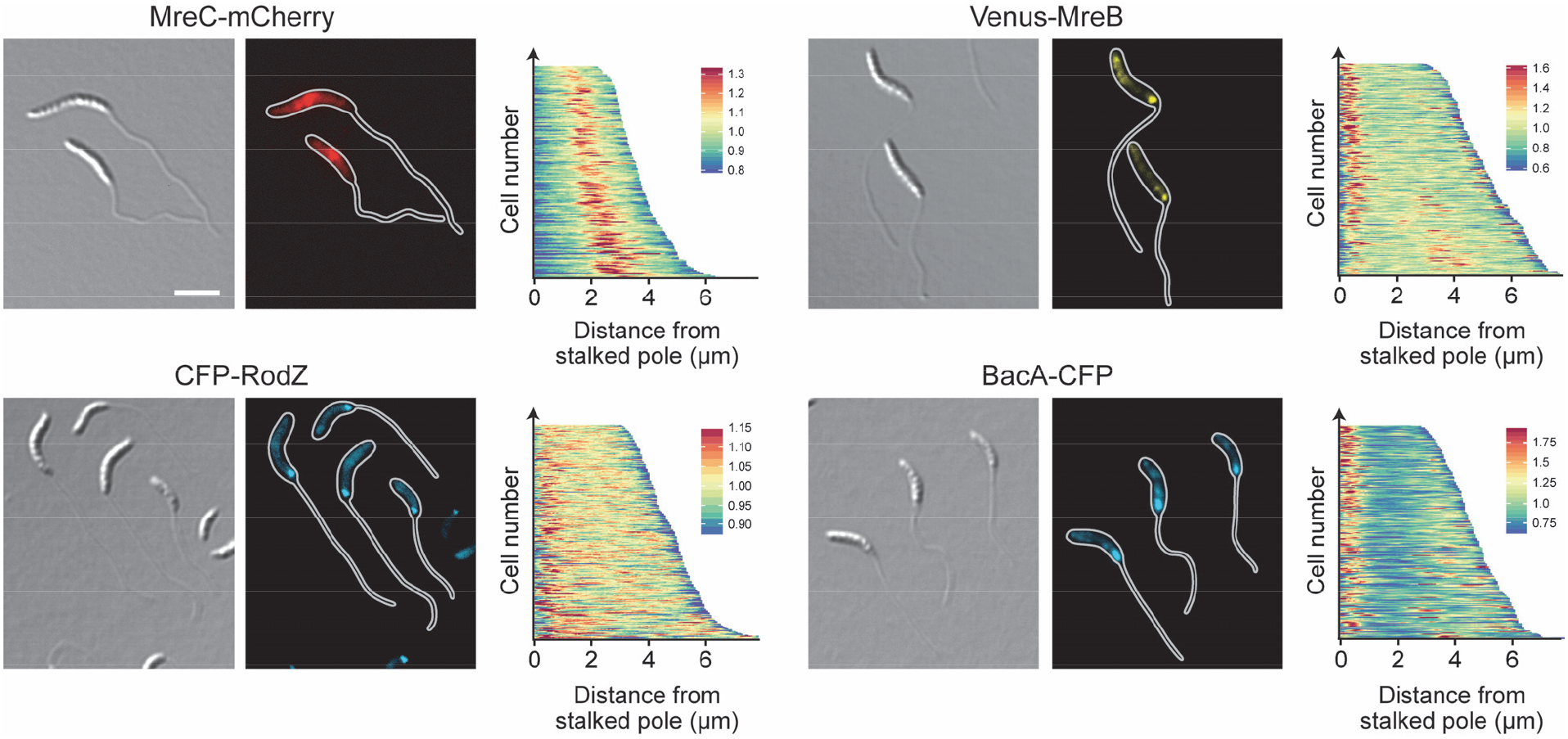
Localization of scaffolding proteins in phosphate-starved cells. Shown is the localization of MreC-mCherry (MAB223, Pxyl::Pxyl-*mreC-mCherry*), CFP-RodZ (CJW2745, *rodZ::cfp-rodZ*), Venus-MreB (MT309, Pxyl::Pxyl-venus-mreB), and BacA-CFP (MT260, *bacA::bacA-cfp*) in cells cultivated for 24 h in M2G^-P^ medium (scale bars: 3 μm). Synthesis of the fluorescent protein fusions was induced with 0.3% xylose 3 h prior to analysis. The population-wide distribution of fluorescence signals was quantified by demographic analysis of random subpopulations of cells (n=200 for each strain).

To determine the role of the different scaffolds in polar PG biosynthesis, cells lacking these factors were subjected to HADA staining after phosphate deprivation (**Figure 10**). Interestingly, despite its severe stalk elongation defect (**Figure 8**) the Δ*bacA* mutant still displayed intense polar foci, indicating that BacA is an accessory factor that is not critical for the global reorganization of PG biosynthesis induced under phosphate-limiting conditions. Consistent with this idea, muropeptide analysis showed that deletion of *bacA* did not have any appreciable effects on global PG composition (**Supplementary file 4**). Depletion of MreB or RodZ, by contrast, strongly decreased the intensity of the polar HADA signals, and frequently led to the insertion of cell wall material at pole-distal sites. In both cases, these defects were accompanied by significant changes in the whole-cell muropeptide profiles. Similar to the Δ*sdpAB* mutant (compare **Supplementary file 2**), the degree of crosslinkage was significantly reduced, mostly due to a decrease in the proportion of highly crosslinked (trimeric and tetrameric) muropeptide species. Moreover, there was a striking increase in the proportion of muropeptides with tripeptide side chains, indicative of high levels of LD-TPase activity. Thus, cell wall stress caused by reduced levels of PBP2-mediated DD-transpeptidation may trigger a fail-safe mechanism that stabilizes the PG meshwork through the formation of abundant 3-3 crosslinks.

**Figure 10.**
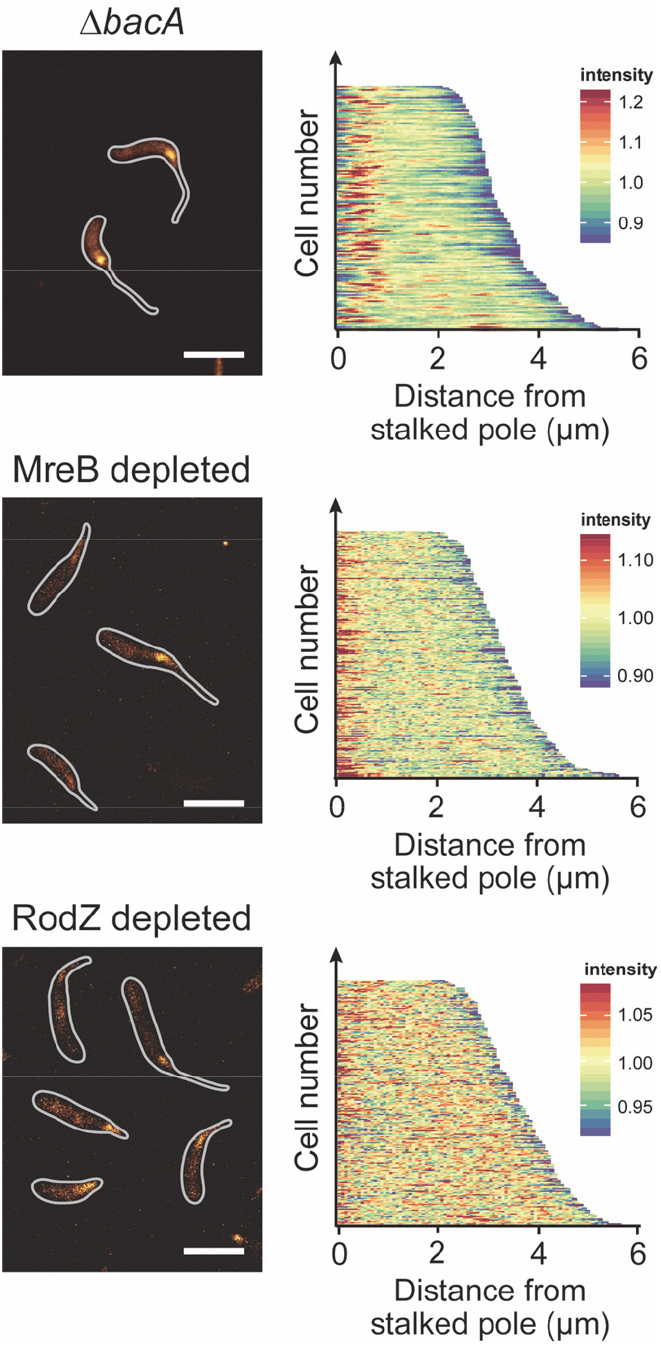
Cell wall biosynthesis in mutants lacking single scaffolding proteins. Shown are fluorescence images of strains LS3809 (Δ*mreB* Pxyl::Pxyl-*mreB*), and CJW2747 (Δ*rodZ::Ω* Pxyl::Pxyl-*rodZ* cultivated as described in **Figure 8A** and subjected to HADA staining (2 min). The population-wide distribution of HADA fluorescence was quantifed by demographic analysis (n=200 for each strain) (scale bars: 3 μm).

Collectively, these results demonstrate that MreB and its transmembrane adapter RodZ play a central role in the establishment of the polar PG biosynthetic zone that gives rise to the stalk structure.

### MreB orchestrates the polar stalk biosynthetic complex

Our data demonstrate that several components of the PG biosynthetic machinery localize to the stalked pole in phosphate-starved cells, suggesting that they assemble into a complex mediating the synthesis of stalk PG. To obtain more insight into the factors mediating the recruitment of these proteins, we reanalyzed the localization patterns of DipM-mCherry, CrbA-mCherry, Venus-MreB, CFP-RodZ, and BacA-CFP in all deletion strains that showed defects in stalk elongation (Δ*dipM*, Δ*sdpAB*, Δ*crbA*, Δ*ldpA*, and Δ*bacA*). However, in all cases, the positioning of the fusion proteins remained unaffected, indicating that neither lytic factors nor the bactofilin cytoskeleton are required for complex assembly. Given the prevalence of elongasome components among the polarly localized proteins, we then tested the role of MreB in the recruitment process. Treatment of cells with the MreB inhibitor A22 not only led to the delocalization of the known MreB interactor RodZ but also abolished the polar foci of DipM and CrbA (**Figure 11**). Thus, MreB appears to be a key organizer of the stalk biosynthetic complex. Notably, A22 had no effect on the polar localization of BacA, indicating that the bactofilin scaffold acts independently of MreB.

**Figure 11.**
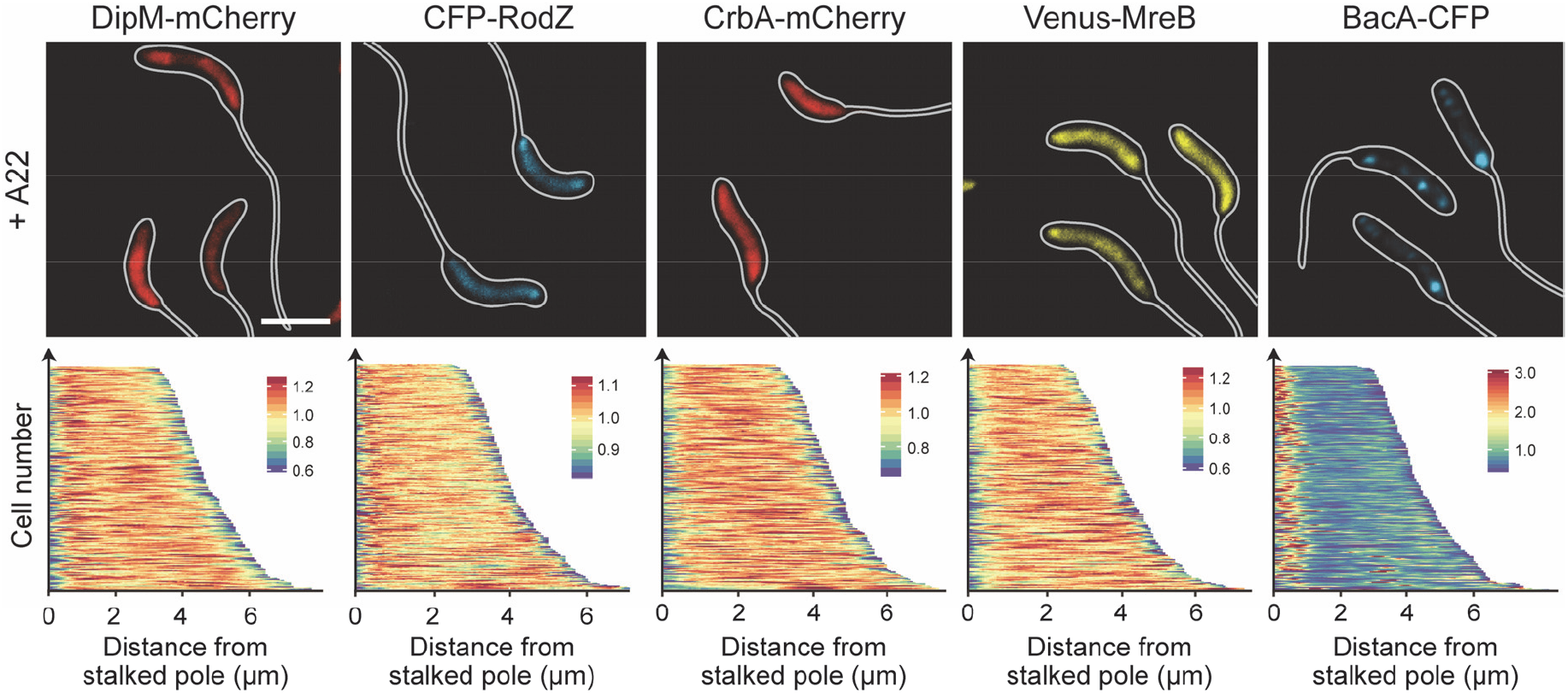
Role of MreB in the polar recruitment of factors involved in stalk formation. Shown are fluorescence images of strains AM208 (Pxyl::Pxyl-*dipM-mCherry*), CJW2745 (*rodZ::cfp-rodZ*), MAB247 (Pxyl::Pxyl-*crbA-mCherry*), MT309 (Pxyl::Pxyl-*venus-mreB*) and MT260 (*bacA::bacA-cfp*). Cells were grown in PYE medium, diluted into M2G^-P^ medium, and incubated for 23 h. Subsequently, A22 (10 μg/ml) was added to the media, and cultivation was continued for 1 h prior to imaging. Strains AM208, MAB247, and MT309 were induced for 3 h with 0.3% xylose to induce synthesis of the fusion proteins before analysis (scale bars: 3 μm).

To analyze the dynamics of the polar MreB assembly, we aimed to construct a sandwich fusion in which mCherry was inserted into a surface-exposed loop of the MreB protein (Bendezu et al., 2009) (**Figure 12A**). A strain carrying the respective allele (*mreB*^sw^) in place of the endogenous *mreB* gene showed normal growth rates (**Figure 12–figure supplement 1A**). However, in rich medium, cells were shorter and more highly curved than the wild type and occasionally formed branches and/or filaments. Under phosphate-limiting conditions, by contrast, the distribution of cell lengths was similar to that of the wild-type strain (**Figure 12–figure supplement 1B**). Strikingly, *mreB*^sw^ cells failed to form stalks under both standard and phosphate-limiting conditions. Consistent with this observation, the fusion protein no longer condensed into polar foci during phosphate starvation but retained the patchy localization pattern typically observed in exponentially growing cells (Gitai, Dye, & Shapiro, 2004) (**Figure 12 C and D**). This unusual behavior led to changes in the global muropeptide profile that were qualitatively similar to those observed for MreB- and RodZ-depleted cells but considerably less pronounced (**Supplementary file 4**). Consistent with the slightly aberrant morphology of the mutant cells, this finding suggests that the insertion of mCherry leads to a mild general defect in MreB function. Importantly, however, it appears to additionally block a specific set of interactions that are critical for the polar recruitment of MreB, thereby preventing stalk formation. To our knowledge this is the first report of a *Caulobacter* strain that is completely stalkless under all growth conditions.

**Figure 12.**
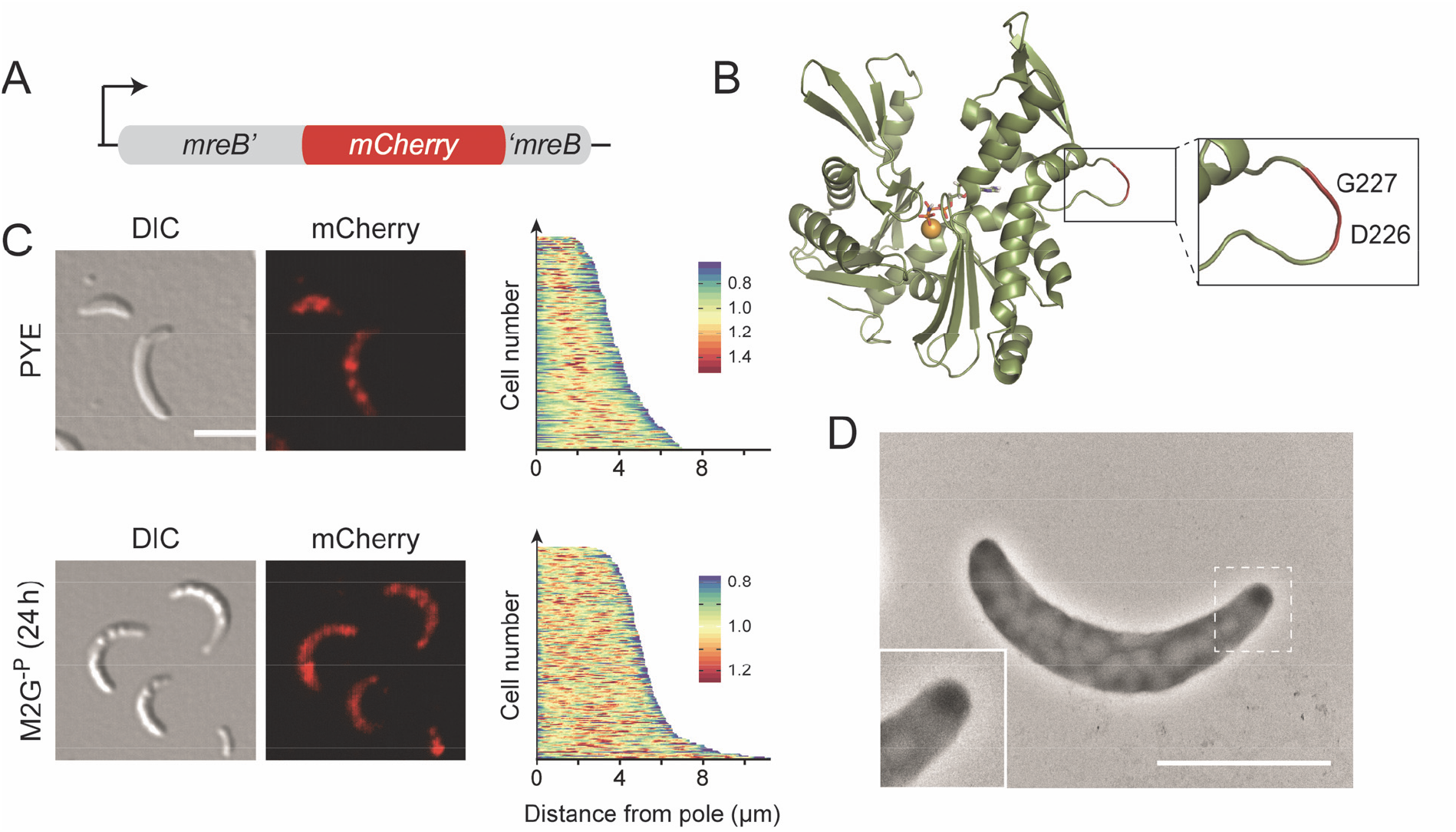
Abolishment of stalk formation in a strain producing an MreB sandwich fusion. (**A**) Schematic representation of the *mreb*^sw^ allele. (**B**) Structure of *Caulobacter* MreB (PDB accession 4CZM; (van den Ent et al., 2014)). The inset shows the site used to insert the mCherry tag. (**C**) DIC and fluorescence images of strain MAB238 (*mreB::mreB^sw^*) grown to exponential phase in PYE medium or incubated for 24 h in M2G^-P^ medium (scale bars: 3 μm). The demographs on the right display the distribution of mCherry fluorescence in random subpopulations of cells (n=210). (**D**) Transmission electron micrograph of MAB238 cells after 24 h of growth in M2G^-P^ medium and staining with uranyl acetate (2%) (scale bar:2 μm).

Collectively, these results demonstrate that MreB has a key role in the assembly and function of the polar stalk biosynthetic complex in *Caulobacter*.

## Discussion

Bacterial cells come in a variety of different shapes, but in most cases the mechanisms generating this morphological diversity are poorly understood (Young, 2006). This study uses the *Caulobacter* stalk as a readily amenable model system to investigate the molecular principles underlying the development of species-specific morphological traits. Previous work has shown that stalk growth is driven by zonal PG incorporation at the old cell pole (Aaron et al., 2007; Kuru et al., 2012; Schmidt and Stanier, 1966). Initially, this process was thought to be mediated by FtsZ and mechanistically similar to pre-septal cell elongation (Divakaruni et al., 2007; Quardokus et al., 1996; Quardokus et al., 2001). However, localization studies revealed that FtsZ is not detectable at the stalked pole, neither during normal cell cycle progression (Thanbichler and Shapiro, 2006) nor during phosphate starvation (**Figure 1B**), excluding the divisome as a relevant player in stalk formation. Other reports implicated MreB and RodZ in stalk growth, suggesting that the elongasome could have a dual role in both cell body and stalk elongation (Divakaruni et al., 2007; Wagner et al., 2005). Clarification of this issue is complicated by the fact that the inactivation of factors with a global role in PG biosynthesis leads to pleiotropic morphological defects. Exploiting the fact that phosphate starvation suppresses *Caulobacter* cell division while strongly promoting stalk elongation, we were able to disentangle cell body- and stalk-specific growth processes and specifically identify proteins involved in the synthesis of stalk PG. Our results indicate that stalk biogenesis is driven by a specialized biosynthetic complex whose composition and biosynthetic activities are clearly distinct from those of the generic cell elongation and division machineries.

Interestingly, the stalk biosynthetic complex is a hybrid composed of factors typically associated with the elongasome (MreB, RodZ, RodA, PBP2) or divisome (DipM, SdpA, SdpB, CrbA) (**Figure 13**). The recruitment of components from the cell elongation machinery may reflect the need to incorporate new cell wall material into an existing sacculus to drive the elongation of the stalk structure, a process that may be mechanistically similar to dispersed PG biosynthesis during lateral cell growth. Notably, HADA (**Figure 2A**) and D-Cysteine (Aaron et al., 2007) labeling clearly indicate that, during stalk growth, newly synthesized PG is not primarily detected in the basal stalk segment but rather in the adjacent polar regions of the cell body. This observation suggests that stalk elongation does not occur simply by addition of new material to the existing stalk template. Instead, it appears to be mediated through expansion of the stalk-proximal polar cap and its simultaneous remodeling into a new stalk segment, a process reminiscent of the medial growth and constriction of the PG sacculus during *Caulobacter* cell division. The common requirement for extensive PG remodeling may explain why the cell division and stalk biosynthetic complexes show a considerable overlap in their autolytic machineries. Interestingly, the importance of some of these shared components varies substantially between the two complexes. For instance, combined inactivation of the lytic transglycosylases SdpA and SdpB has no obvious effect on cell division (Zielinska et al., 2017), whereas it largely abolishes stalk formation, indicating that the functional context of these proteins varies depending on the process they mediate. It remains to be clarified to what extent the different cell wall biosynthetic complexes compete for their shared components. Interestingly, during the *Caulobacter* cell cycle, stalk growth occurs predominantly within a short time window at the transition from dispersed to medial peptidoglycan biosynthesis (**Figure 2–figure supplement 1**). It is therefore tempting to speculate that the elongasome and divisome have higher priority in the recruitment of shared factors, thereby restricting assembly of the stalk biosynthetic complex to phases in which they are not fully active. Overall, stalk formation clearly demonstrates how the reshuffling of preexisting machinery can serve as a straightforward means to generate novel morphological features in bacteria. The striking diversity of cell shapes observed in certain lineages, such as the alphaproteobacteria, may therefore not be based on major new additions to the repertoire of cell wall biosynthetic proteins but rather on subtle changes in protein activities and localization patterns.

**Figure 13.**
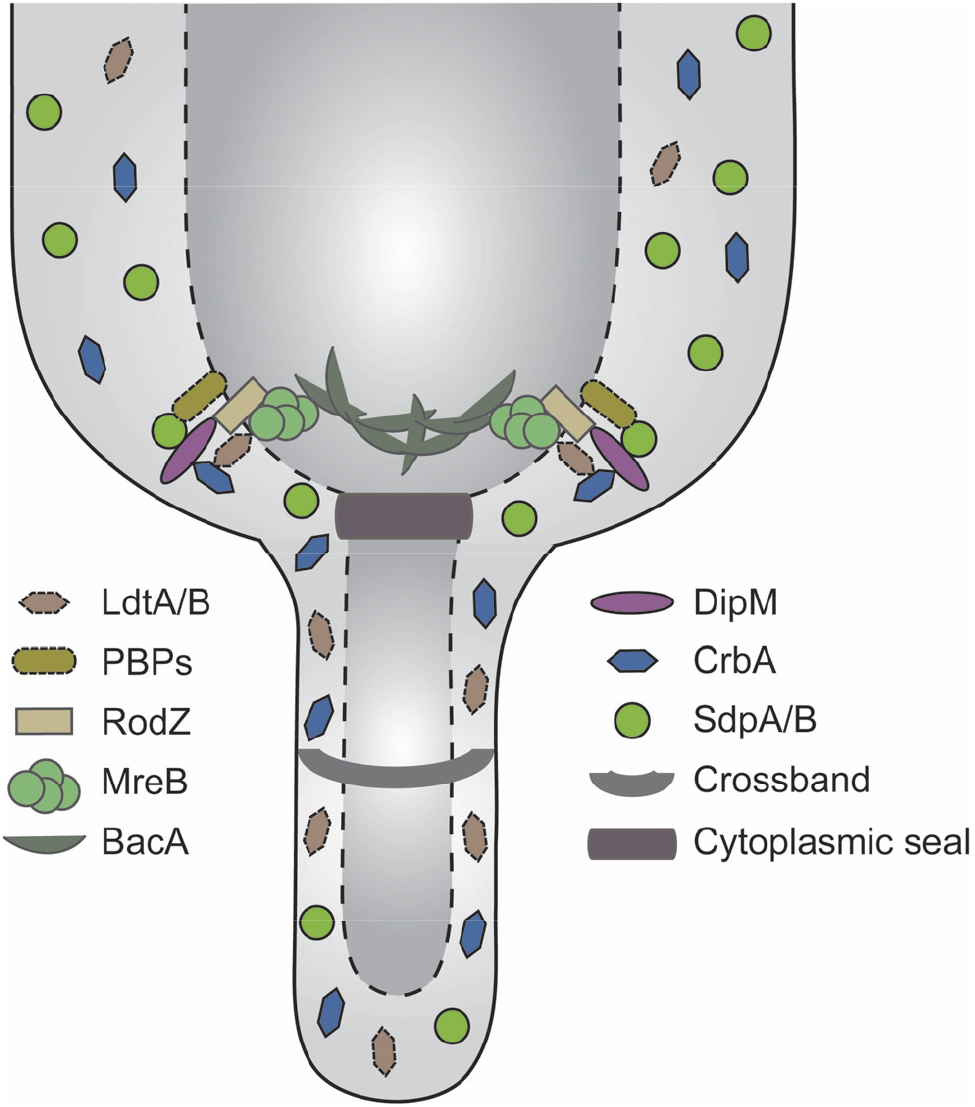
Model of the factors contributing to stalk formation. The actin homolog MreB forms the basis of the stalk biosynthetic complex and mediates the recruitment of both synthetic and lytic proteins to the stalked pole. The bactofilin BacA contributes to stalk biosynthesis but is not essential to this process. In addition to the proteins that are stably associated with the cell pole, diffusible periplasmic enzymes are transiently associated with the polar machinery or non-selectively trapped in the compartments generated by the crossbands.

A key finding of our work is the central role of MreB in the stalk biosynthetic complex. We show that this cytoskeletal protein condenses at the stalked pole during phosphate starvation and facilitates the polar recruitment of several other factors that are critical to stalk formation. Notably, our attempts to integrate mCherry into a surface-exposed loop of MreB led to the serendipitous identification of a *Caulobacter* strain that was completely devoid of stalks under both phosphate-limiting and -replete conditions. This is in stark contrast to other mutants described previously, which are stalk-less in rich medium but still elaborate stalks upon phosphate starvation (Biondi et al., 2006; Bowman et al., 2008; Ebersbach et al., 2008; Sommer and Newton, 1989), suggesting that they have a defect in the regulation of stalk formation rather than in the biosynthetic machinery mediating this process. Importantly, cells producing the MreB sandwich fusion showed only relatively mild general cell shape defects. The region surrounding the insertion site of mCherry may thus contain determinants that are specifically required for MreB’s function in stalk formation but largely dispensable for elongasome-mediated longitudinal growth of the cell body. Previous work has shown that the positioning of MreB filaments is strongly influenced by their intrinsic curvature (Hussain et al., 2018; Ursell et al., 2014), a parameter controlled by the concentration of the membrane adapter RodZ (Colavin et al., 2018). The high enrichment of the MreB-RodZ complex at the stalked pole may thus be sufficient to change the architecture of MreB filaments such as to faciliate their interaction with the more highly curved stalked pole. However, the cues promoting the relocation of MreB from the lateral regions of the cell to the stalked pole still remain unknown.

Although MreB clearly has a key role in stalk biogenesis, it is not the only scaffolding protein contributing to this process. Previous work has shown that the bactofilin BacA is required for proper stalk length {Kühn, 2010 #41}, and our analyses revealed an additional role for this protein in cell body elongation during phosphate starvation (**Figure 8**). Notably, the *bacA* gene lies in a putative operon with *ldpA*, a gene encoding a putative LytM-like endopeptidase that also functions in stalk formation. This genetic context is conserved in a variety of other species, suggesting a functional link between the two gene products (Jackson et al., 2018; Sycuro et al., 2010). Support for this notion comes from studies in the human pathogen *Helicobacter pylori*, which demonstrated that both genes in this conserved operon are required to establish the characteristic helical cell shape of this species (Sycuro et al., 2010). Notably, apart from its putative interaction with LdpA, *Caulobacter* BacA was shown to recruit a class A PBP (PbpC) involved in stalk elongation and in the targeting of proteins to the stalk lumen (Hughes et al., 2013; Kühn et al., 2010). Importantly, the polar localization of BacA was independent of the presence of MreB. The bactofilin cytoskeleton thus appears to constitute a functionally independent morphogenetic module that has been coopted by *Caulobacter* to modulate stalk formation. This module appears to act downstream of the MreB-dependent stalk biosynthetic complex, as it was not able to establish a stalk structure in the absence of a functional MreB cytoskeleton.

The ultimate determinant mediating the polar recruitment of the stalk biosynthetic machinery in *Caulobacter* still remains unknown. In *Asticcacaulis excentricus*, a member of the *Caulobacteraceae* that is characterized by subpolar stalks, the site of stalk formation was shown to be defined by the polarity determinant SpmX (Jiang et al., 2014). However, despite its conservation, this protein is not required for proper stalk localization in *Caulobacter* cells (Radhakrishnan et al., 2008). Notably, deletion of SpmX or transfer of the cells to phosphate-limited media restores polar stalk growth in *A. excentricus* (Jiang et al., 2014), suggesting that the pathway observed in *Caulobacter* is still present in this species but normally obscured by the the action of the newly coopted localization factor SpmX. It will be interesting to see whether the *A. excentricus* SpmX homolog organizes an alternative stalk biosynthetic complex or simply recruits the polar machinery to a pole-distal position.

Although the functionality and localization of the peptidoglycan biosynthetic machinery changes drastically upon transition of *Caulobacter* cells from phosphate-replete to phosphate-limiting media, the overall composition of their PG layer remains largely unaffected. This finding is unexpected because significant changes in both glycan chain lengths and the degree of cross-linking were observed in other species in response to changes in their growth conditions (Vollmer et al., 2008). However, analyzing the muropeptide profiles of isolated stalk and cell body fractions, we identified clear differences between these two compartments that are likely obscured in whole-cell analyses due to the small contribution of stalks to the total cellular PG content. Most importantly, stalk PG showed a significantly higher degree of crosslinkage, which was mostly due to a higher frequency of 3-3 crosslinks, indicative of elevated LD-TPase activity. The precise reason for this difference remains unclear. It is conceivable that the LD-TPases LdtD and LdtX are part of the polar stalk biosynthetic complex and, thus, preferentially act on newly synthesized PG produced by this machinery. However, localization studies did not give any evidence for an enrichment of these proteins at the stalked pole. An alternative explanation may be provided by the observation that the turnover rate of PG is significantly lower in the stalk than in the cell body. Thus, LD-TPases may act uniformly throughout the entire cell envelope, but most of the 3-3 crosslinks formed in the cell body may be lost as a consequence of PG remodeling, whereas those in the stalk are retained over prolonged periods of time. Notably, peptides with 3-3 crosslinks are stiffer than those with 3-4 crosslinks and adopt a more extended conformation that is better suited to connect glycan strands in stressed PG (de Pedro and Cava, 2015). Their increased frequency may therefore help to modulate the mechanical properties of the stalk and render it more resistant to bending or breakage under conditions of high laminar flow (Klein et al., 2013; Persat et al., 2014)

Collectively, our study shows that, in *Caulobacter*, multiple cell-wall biosynthetic machineries act in concert to generate stalks of proper size and stability, thereby ensuring optimal performance of this cellular structure in the environmental context. It will be interesting to see how the nature and the regulation of these components have changed during evolution to bring about the large variety of morphologies found in other stalked members of the alphaproteobacterial lineage.

## Materials and Methods

### Media and growth conditions

*Caulobacter* strains (Evinger and Agabian, 1977) were grown at 28°C in peptone-yeast-extract (PYE) medium (Poindexter, 1964), supplemented with antibiotics at the following concentration when appropriate (μg ml^-1^; liquid/solid medium): spectinomycin (25/50), streptomycin (-/5), gentamicin (0.5/5), kanamycin (5/25), chloramphenicol (1/1). Gene expression from the *xylX* promoter (Pxyl) or *vanA* promoter (Pvan), was induced by supplementation of the media with 0.3% D-xylose and 0.5 mM sodium vanillate, respectively, prior to analysis of the cells. To induce phosphate starvation, stationary cells were diluted 1:20 in M2G^-P^ medium (Kühn et al., 2010) and incubated at 28°C for the indicated times. In case of the conditional *mreB, rodZ*, and *mreC* mutants, cells were grown to exponential phase (OD_600_ ^~^ 0.5) in PYE medium supplemented with xylose, washed three times, and then resuspended to an OD_600_ of 0.05 in inducer-free medium. The cultures were then grown for 7 h to achieve protein depletion, diluted (1:20) in M2G^-P^ medium, and cultivated for additional 24 h before analysis. The conditional *amiC* and *dipM* mutants were treated in a similar fashion, with 12 h of cultivation in PYE medium prior to transfer into M2G^-P^. The synchronization of *Caulobacter* was achieved by density gradient centrifugation using Percoll (Sigma-Aldrich) (Tsai and Alley, 2001). To determine the viable-cell count in cultures, various dilutions of the cell suspensions were spread on PYE plates, and the number of colony-forming units (CFU) was determined after three days of incubation at 28 °C. *E. coli* strain TOP10 (Invitrogen) and its derivatives were cultivated at 37°C in LB broth (Karl Roth, Germany). Antibiotics were added at the following concentrations (μg/ml; liquid/solid medium): spectinomycin (50/100), gentamicin (15/20), kanamycin (30/50), chloramphenicol (20/30).

### Plasmid and strain construction

The bacterial strains, plasmids, and oligonucleotides used in this study are listed in Supplementary file 5. *E. coli* TOP10 (Invitrogen) was used as host for cloning purposes. All plasmids were verified by DNA sequencing. *Caulobacter* was transformed by electroporation. Non-replicating plasmids were integrated into the *Caulobacter* chromosome by single-homologous recombination at the *xylX* (Pxyl) or *vanA* (Pvan) locus (Thanbichler et al., 2007). Gene replacement was achieved by double-homologous recombination using the counter-selectable *sacB* marker (M.R.K. Alley, unpublished) (Thanbichler & Shapiro, 2006). Proper chromosomal integration or gene replacement was verified by colony PCR.

### Growth curves

Cells were grown to exponential phase in PYE medium, harvested by centrifugation, and resuspended in the same medium to an OD_600_ of 0.05. The suspensions were then transferred to 24-well polystyrene microtiter plates (Becton Dickinson Labware), incubated at 32°C with double-orbital shaking in an Epoch 2 microplate reader (BioTek, Germany), and analyzed photometrically (OD_600_) at 15 min intervals.

### Light and immunofluorescence microscopy

For light microscopic analysis, cells were transferred onto pads made of 1% agarose. Images were taken with an Axio Observer.Z1 (Zeiss) microscope equipped with a Plan Apochromat 100x/1.45 Oil DIC and a Plan Apochromat 100x/1.4 Oil Ph3 phase contrast objective, an ET-mCherry filter set (Chroma, USA), and a pco.edge sCMOS camera (PCO). Images were recorded with VisiView 3.3.0.6 (Visitron Systems, Germany) and processed with Metamorph 7.7.5 (Universal Imaging Group, USA) and Illustrator CS6 (Adobe Systems, USA). To generate demographs, fluorescence intensity profiles were measured with ImageJ 1.47v (http://imagej.nih.gov/ij). The data were then processed in R version 3.5.0 (Team, 2012) using the Cell Profiles script (http://github.com/ta-cameron/Cell-Profiles) (Cameron et al., 2014). Box and violin plots for the statistical analysis of imaging data were generated in R version 3.5.0 using the ggplot2 (Wickham, 2009) and Reshape2 (Wickham, 2007) packages, respectively.

### Electron microscopy

10 μl cell suspension were applied to an electron microscopy grid (Formvar/Carbon Film on 300 Mesh Copper; Plano GmbH, Germany) and incubated for 1 min at room temperature. Excess liquid was removed with Whatman filter paper. Subsequently, the cells were negatively stained for 5 sec with 5 μl of 1% uracyl acetate. After three washes with H_2_O, the grids were dried, stored in an appropriate grid holder, and analyzed in a 100 kV JEM-1400 Plus transmission electron microscope (JEOL, USA).

### Western blot analysis

Western blot analysis was performed as described (Thanbichler and Shapiro, 2006), using anti-CtrA (Domian et al., 1997), anti-FtsZ (Goley et al., 2010), anti-MipZ (Thanbichler and Shapiro, 2006), anti-DnaA (Collier et al., 2006), or anti-SpmX (Radhakrishnan et al., 2008) at dilutions of 1:10,000 (anti-CtrA, anti-FtsZ, anti-MipZ, and anti-DnaA), and 1:50,000 (anti-SpmX). Goat anti-rabbit immunoglobulin G conjugated to horseradish peroxidase (Perkin Elmer, USA) was used as secondary antibody. Immunocomplexes were detected using the Western Lightning Plus-ECL chemilumines-cence reagent (Perkin Elmer, USA). Signals were recorded with a ChemiDoc MP imaging system (Bio-Rad) and analyzed using the Image Lab 5.0 software (Bio-Rad).

### HADA staining

HADA-staining experiments were conducted as described (Kuru et al., 2012). Briefly, 50 μl of a culture were incubated for 2 min with 0.5 mM HADA. The cells were then fixed by addition of ice-cold ethanol to a concentration of 70% and incubated at 4°C for 20 min. Subsequently, they were washed three times with PBS and subjected to fluorescence microscopic analysis. For chase experiments, phosphate-starved *Caulobacter* cells were grown for 90 min in the presence of 0.5 mM HADA. The cells were washed three times with M2G^-P^ medium, resuspended in fresh M2G^-P^ medium, and further cultivated for the indicated time intervals. Cells were fixed and washed as described above prior to imaging.

### Bioinformatic analysis

Protein sequences containing the indicated domains were retrieved from the UniProt Knowledgebase (The Uniprot Consortium, 2017). Their overall domain composition was determined using the SMART server (Letunic et al., 2015). The prediction of protein localization and membrane topology was performed with Signal-BLAST (Frank and Sippl, 2008) and TMHMM (Krogh et al., 2001), respectively.

### Flow cytometry

Cultures were grown in the indicated media and supplemented with 20 μg/ml rifampicin 3 h prior to analysis to block the re-initiation of chromosome replication. At the indicated time points, cells were diluted to an OD_600_ of 0.1-0.2, incubated for 25 min under vigorous shaking with the DNA-specific fluorescent dye Hoechst 33342 (10 μM; ThermoFischer, Germany), and fixed by addition of ethanol to a final concentration of 70%. Subsequently, the suspensions were analyzed by flow cytometry in a customized Fortessa Flow Cytometer (BD Biosciences), using the UV 440/40 nm channel. Data were acquired with FACSdiva 8.0 (BD Biosciences) and processed with FlowJo v10 (FlowJo LLC).

### Peptidoglycan analysis

For whole-cell analyses, cultures were rapidly cooled to 4 °C and harvested by centrifugation at 16,000 rpm for 30 min. The cells were resuspended in 6 ml of ice-cold H2O and added dropwise to 6 ml of a boiling solution of 8% sodium dodecylsulfate (SDS) that was stirred vigorously. After 30 min of boiling, the suspension was cooled to room temperature. Peptidoglycan was isolated from the cell lysates as described previously (Glauner, 1988) and digested with the muramidase cellosyl (kindly provided by Hoechst, Frankfurt, Germany). The resulting muropeptides were reduced with sodium borohydride and separated by HPLC following an established protocol (Bui et al., 2009; Glauner, 1988). The identity of eluted fragments was assigned based on the retention times of known muropeptides from *Caulobacter* (Takacs et al., 2013).

To prepare stalk and cell body fractions, 100 ml cultures grown in M2G^-P^ medium were rapidly cooled to 4 °C and harvested by centrifugation at 16,000 rpm for 30 min. After resuspension in M2G^-P^ medium, the cells were vigorously agitated for 2 min at maximum speed in a kitchen blender. The suspension was submitted to three rounds of centrifugation at 9,000 rpm and 4 °C. The supernatants (stalk fraction) and the first pellet (cell body fraction) were collected separately and kept in ice. The stalk fraction was subjected to an additional centrifugation step at 10,000 rpm and 4 °C to remove residual cell bodies and cell debris. Subsequently, stalks were collected by centrifugation at 20,000 rpm and 4 °C for 30 min, resuspended in 3 ml ice-cold H_2_O, added dropwise to 3 ml of a boiling 8% SDS solution, and then further processed as described above to isolate stalk PG. The isolation of cell body PG was achieved as described for whole-cell samples.

## Acknowledgements

We thank Julia Rosum (University of Marburg) and Lisa Atkinson (Newcastle University) for excellent technical assistance. Moreover, we acknowledge Andrea Möll and Aleksandra Zielinska for support in the initial phases of this work and Manuel Osorio Valeriano and Maria Perez Burgos for help with the transmission electron microscopic analyses. This work was supported by intramural funds from Philipps-Universität Marburg (to M.T.), a Max Planck Fellowship from the Max Planck Society (to M.T.), a Young Investigator Grant (RGY0076/2013-C104) from the Human Frontier Science Program (to M.T.), a grant from the Wellcome Trust (101824/Z/13/Z; to W.V.), and funds from the German Research Foundation (DFG) granted in the context of the Collaborative Research Center *“Microbial Diversity in Environmental Signal Response”* (SFB 987; to M.T.).

---

**Figure 2–figure supplement 1.**
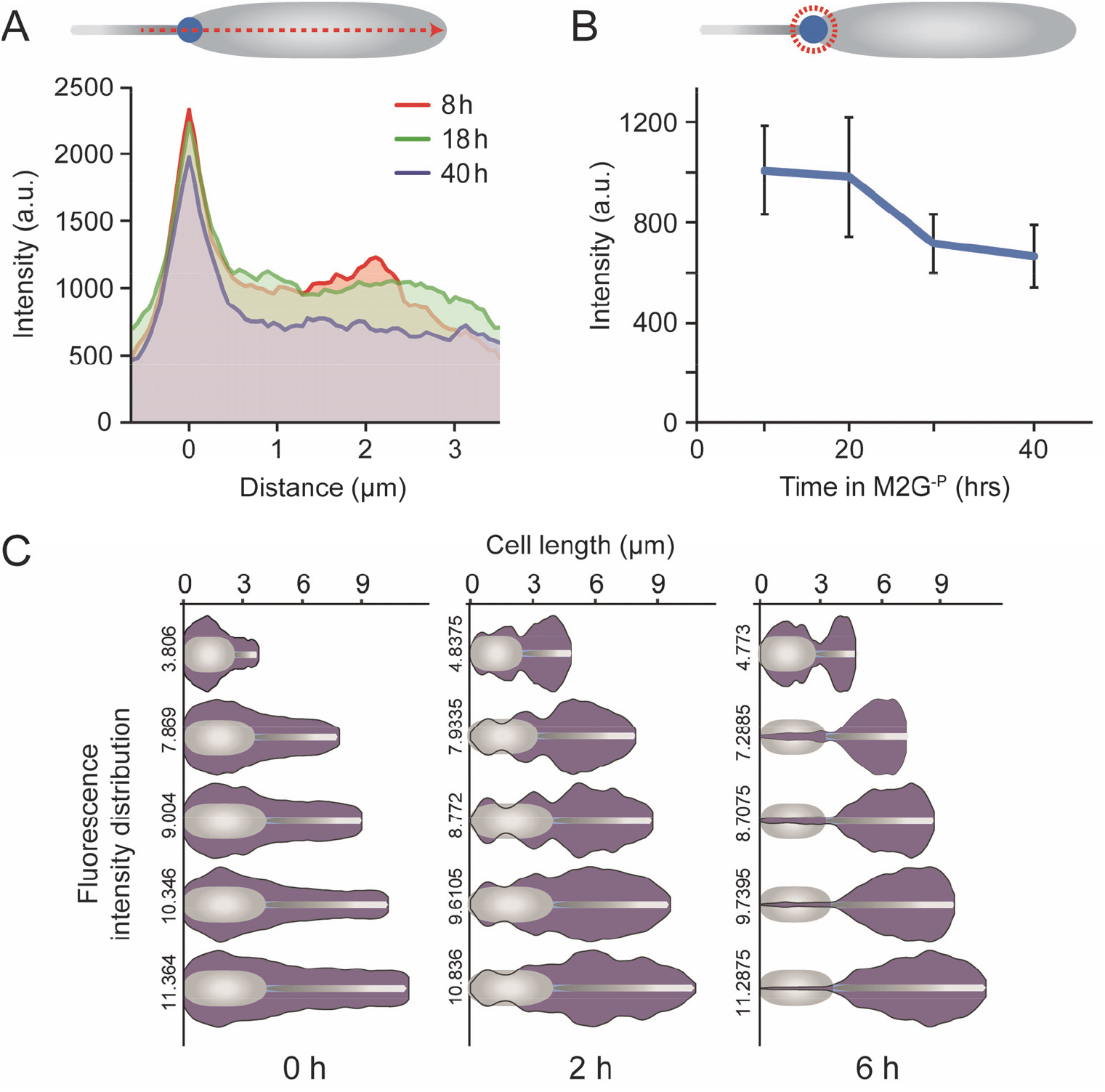
HADA incorporation in wild-type cells in the presence of phosphate. Wild-type (NA1000) swarmer cells were transferred into PYE medium and cultivated for the duration of one cell cycle. At the indicated time points, samples were taken, pulse-labeled (2 min) with HADA, and subjected to fluorescence microscopy (scale bar: 3 μm). The demographs show the distribution of HADA fluorescence in random subpopulations of cells (n=200 per time point).

**Figure 2–figure supplement 2.**
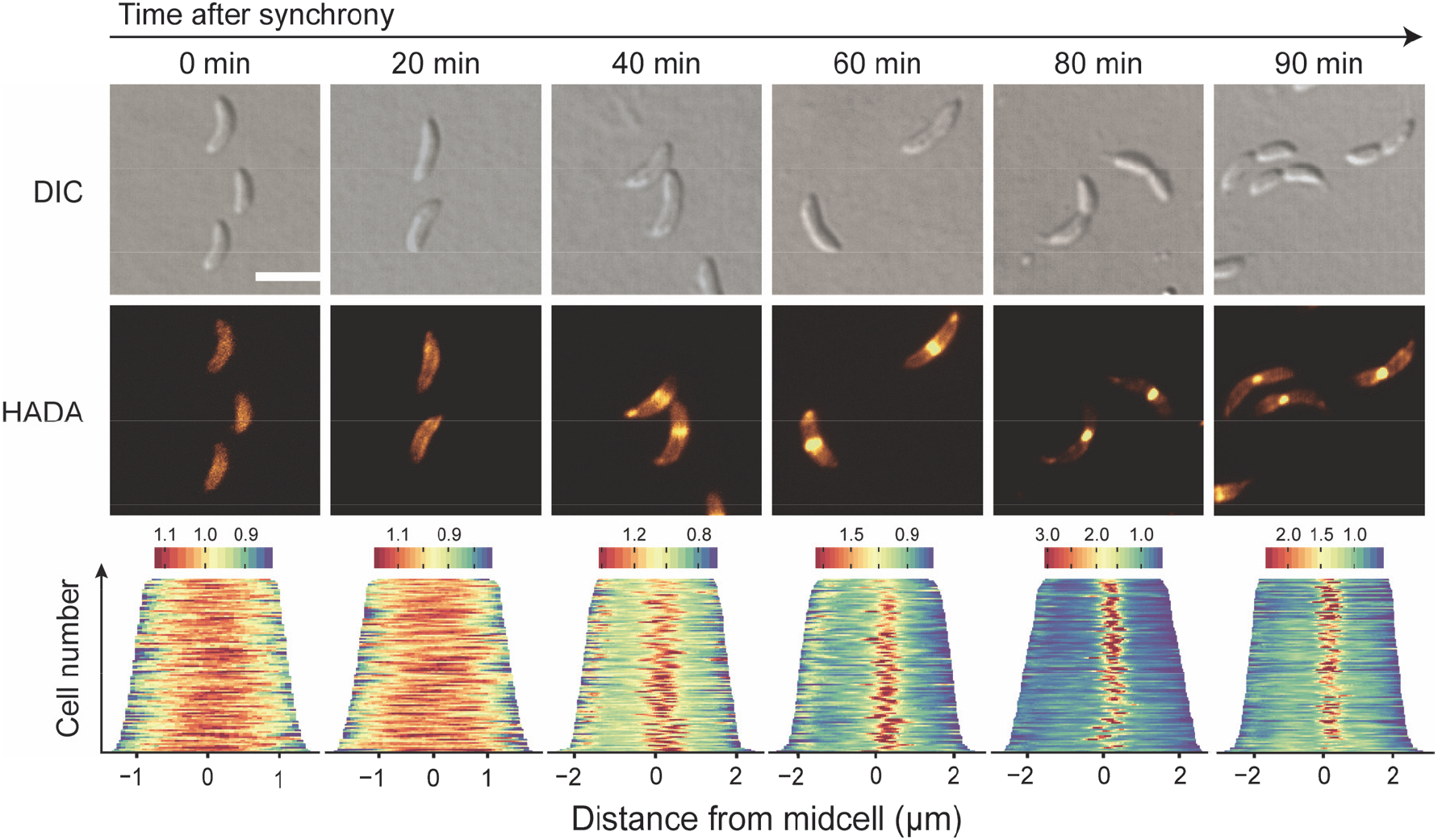
Changes in HADA incorporation during transition to phosphate starvation. (**A**) Distribution of newly synthesized PG after different times of phosphate starvation. Cells of wild-type strain NA1000 were cultiveated in M2G^-P^ medium for the indicated amount of time and exposed to a short (2 min) pulse of HADA. After microscopic analysis, the distribution of fluorescence along the long axis of the cells was determined by line scan analysis for multiple cells per time point. The curves obtained were normalized to the average cell length of the population analyzed, aligned at the center of the stalked-pole focus and averaged (n=42 at 8 h, n=40 at 18 h, and n=44 at 40 h). (**B**) Intensity of HADA fluorescence at the stalked pole in wild-type (NA1000) cells cultivated in M2G^-P^ medium for 8 h (n=51), 18 h (n=60), 28 h (n=54), and 40 h (n=54). Error bars represent standard deviations. (**C**) Slow turnover of PG in the stalk. Cells were cultivated in M2G^-P^ medium for 18 h and exposed to HADA for an extended period of time (1.5 h) to uniformly label their peptidoglycan layer. Subsequently, they were washed, transferred into HADA-free M2G^-P^ medium, and cultivated for 2 h, 4 h, and 6 h in the absence of the label (scale bars: 3 μm). To quantify the changes in HADA fluorescence overtime, fluorescence profiles were obtained from random subpopulations of cells (n=200 per time point). The lengths of the profiles in each quintile of the cell length distribution were normalized to the maximum cell length in the respective quintile, and the fluorescence intensities were averaged and shown as violin plots.

**Figure 3–figure supplement 1.**
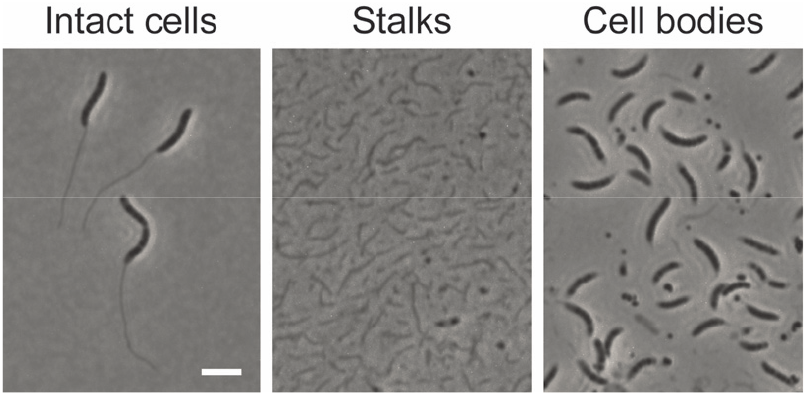
Visualization of isolated stalk and cell body fractions. Cells were cultivated for 24 h in M2G^-P^ medium, agitated vigorously, and then subjected to differential centrifugation to separate stalks and cell bodies. Samples of the intact cells and the stalk and cell body fractions were visualized by phase contrast microscopy (scale bar: 3 μm).

**Figure 4–figure supplement 1.**
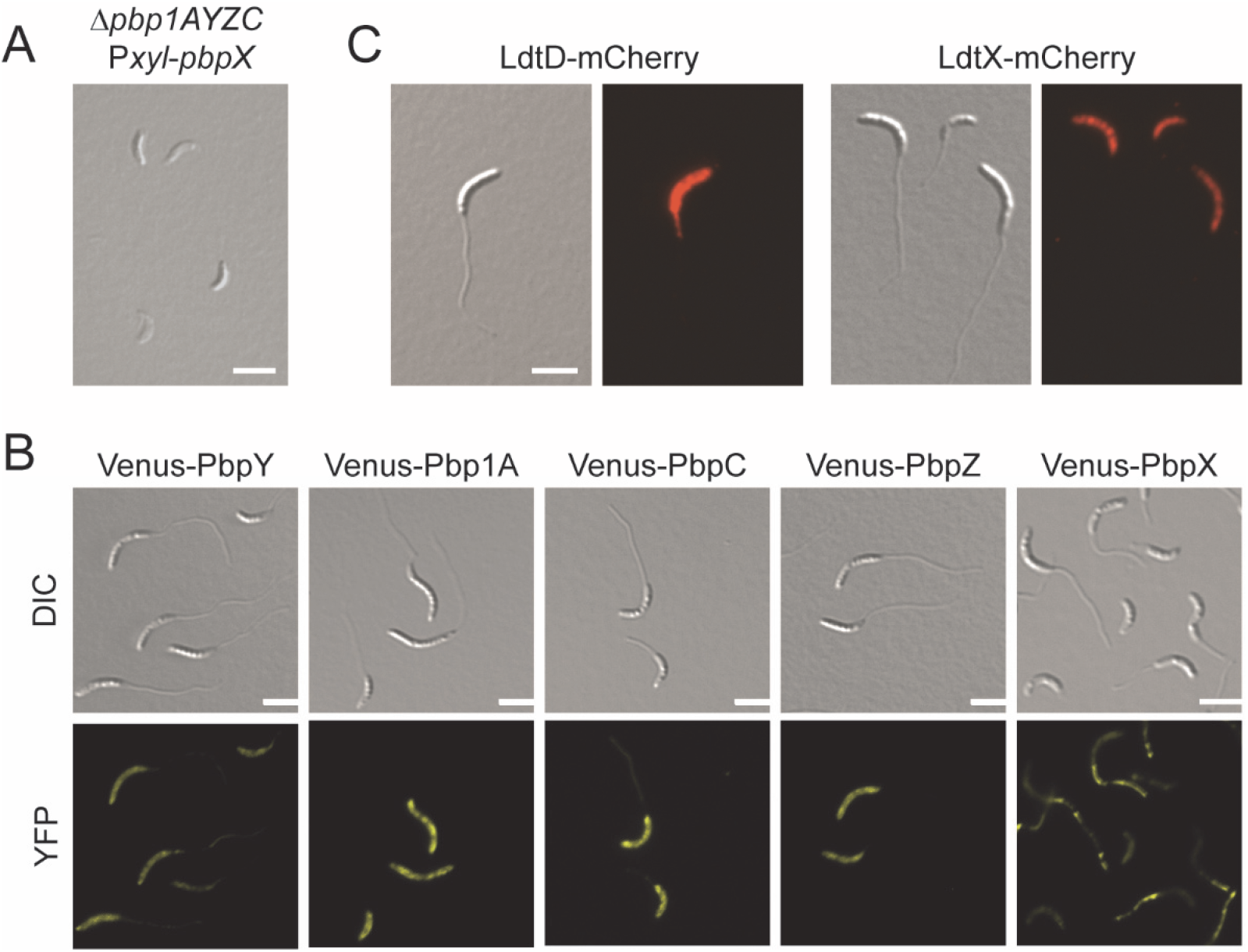
Role of PG synthases in stalk elongation under phosphate starvation. (**A**) Lysis of a conditional mutant lacking all class A PBPs during cultivation in phosphate-limiting conditions. Cells of strain WS056 (Δ*pbpY* Δ*pbp1A* Δ*pbpC* Δ*pbpZ* Pxyl::Pxyl-*pbpX*) were pre-grown until exponential phase in PYE medium containing the inducer xylose, transferred in PYE without xylose after two washing steps, and grown for additional 12 h until they reached stationary phase. Then cells were diluted (1:20) into xylose-free M2G^-P^ medium, and cultivated for 24 h prior to visualization by DIC microscopy (scale bar: 3 μm). (**B**) Localization of fluorescently tagged class A PBPs under conditions of phosphate starvation. Cells of strains AM457 (Pxyl::Pxyl-*venus-pbpY*), KK33 (Pxyl::Pxyl-*venus-pbp1a*), MT279 (Pxyl::Pxyl-*venus-pbpC*), AM458 (Pxyl::Pxyl-*venus-pbpZ*), and MT278 (Pxyl::Pxyl-*venus-pbpX*), were grown for 24 h in M2G^-P^ medium and visualized by fluorescence microscopy. Three hours prior to analysis, the cultures were supplemented with 0.3% xylose to induce synthesis of the fusion proteins (scale bars: 3 μm). (**C**) Localization of fluorescently tagged LD-TPases under conditions of phosphate starvation. Shown are cells of strains MAB389 (Pxyl::Pxyl-*ldtD-mCherry*) and MAB390 (Pxyl::Pxyl-*ldtX-mCherry*) cutivated and induced as described for panel B (scale bar: 3 μm).

**Figure 5–figure supplement 1.**
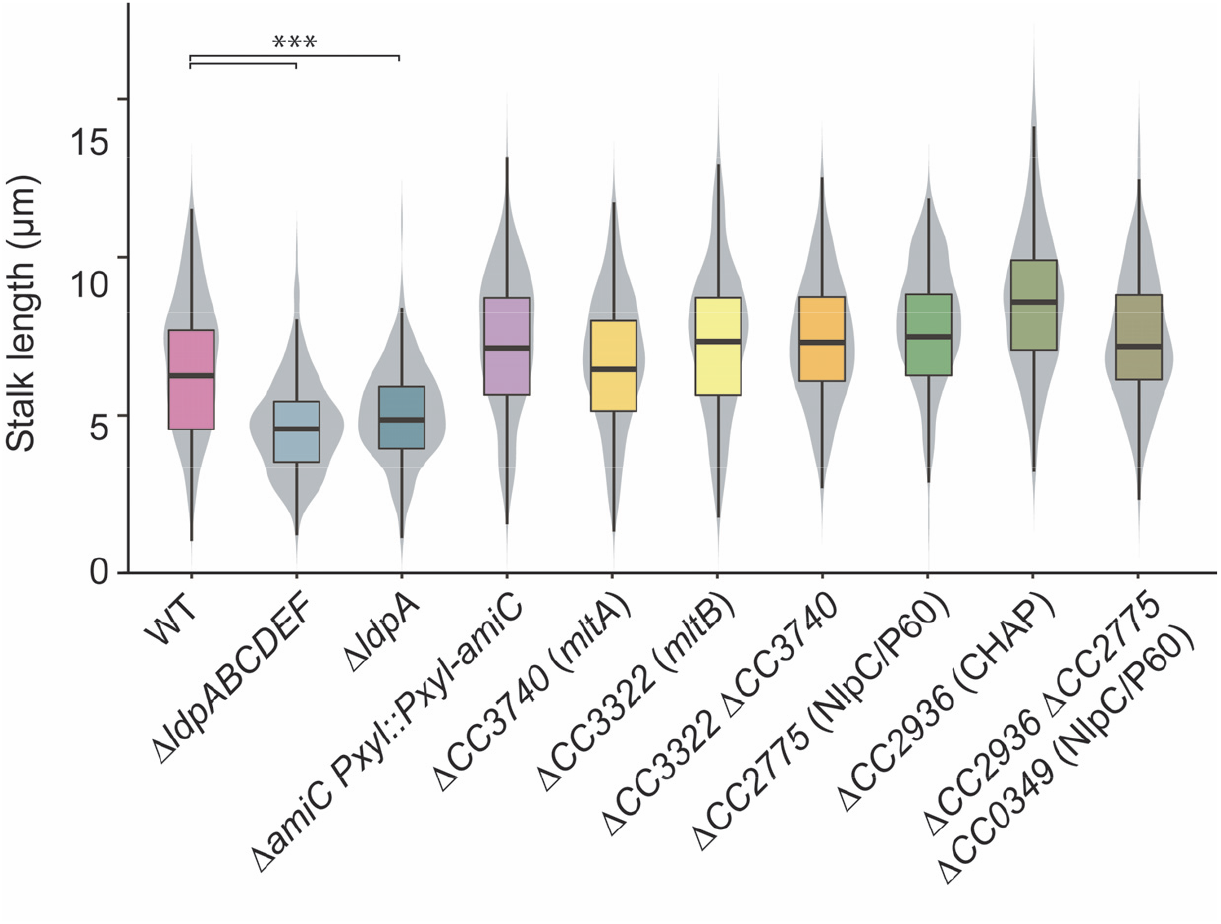
Role of autolytic enzymes in stalk elongation under phosphate starvation. (**A**) Distribution of stalk lengths in populations of mutants lacking predicted autolytic enzymes. Shown are cells of strains AZ52 (Δ*ldpABCDEF*), AM364 (Δ*ldpA*), MAB386 (Δ*amiC* Pxyl::Pxyl-*amiC*), MAB239 (Δ*CC3740*), MAB233 (ΔCC3322), MAB251 (*ΔCC3322 ΔCC3740*), AZ85 (Δ*CC2775*), MAB248 (Δ*CC2936*), and MAB250 (Δ*CC2936* Δ*CC2775* Δ*CC0349*) harvested after 24 h of cultivation in M2G^-P^ medium. The values obtained (n=210 per strain) are shown as box plots, with the thick line indicating the median, the box the interquartile range and the wiskers the 2^nd^ and the 98^th^ percentile. In addition rotated kernel density plots (grey) are depicted for each dataset to indicate the distribution of the raw data (*** p < 10^-6^; t-test).

**Figure 5–figure supplement 2.**
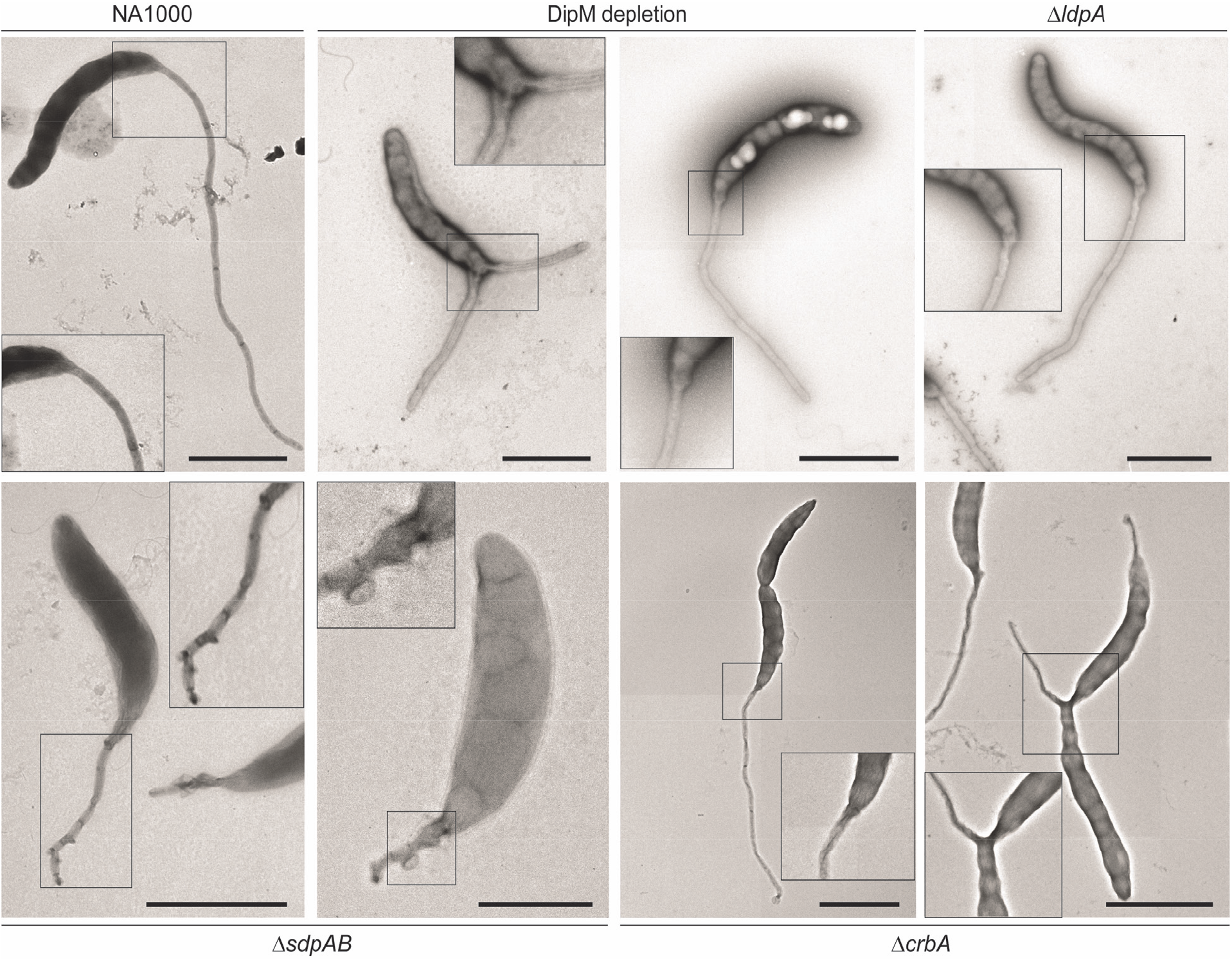
Transmission electron micrographs of mutants lacking autolytic enzymes. Cells of strains NA1000 (WT), MAB360 (Δ*dipM* Pxyl::Pxyl-dipM), AM364 (Δ*ldpA*), AZ22 (Δ*sdpAB*), and AM376 (Δ*crbA*) were grown for 24 h in M2G^-P^ medium, stained with uranyl acetate, and visualized by transmission electron microscopy (scale bars: 2 μm).

**Figure 6–figure supplement 1.**
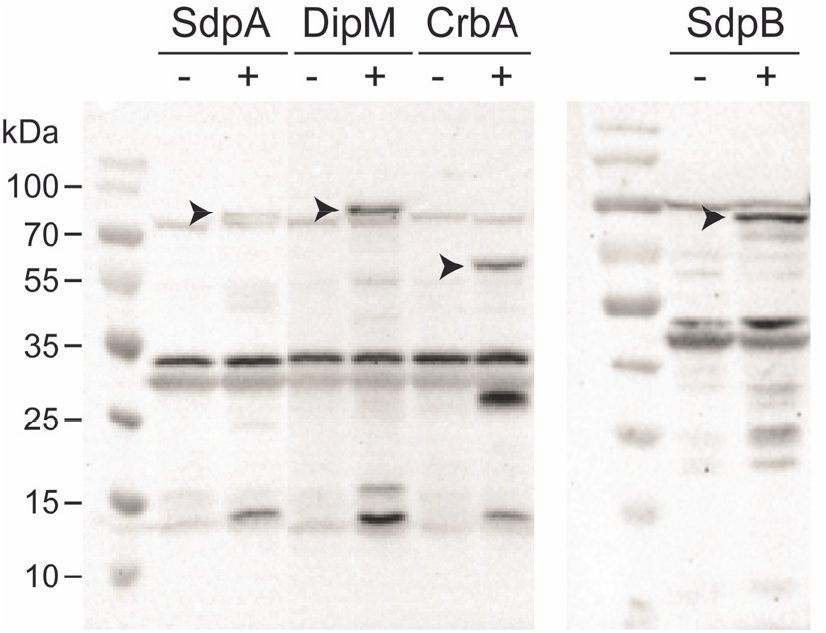
Stability of fluorescent protein fusions. Western blot analysis of strains producing fluorescently tagged derivatives of SdpA (AM480, Pxyl::Pxyl-*sdpA-mCherry*), DipM (AM208, Pxyl::Pxyl-*dipM-mCherry*), CrbA (MAB247, Pxyl::*Pxyl-crbA-mCherry*), and SdpB (AZ127, Pxyl::Pxyl-*torA’-sdpB-mCherry*). Cells were grown for 24 h in M2G^-P^ medium and subjected to microscopy. Three hours (AM480, MAB247, and AM208) or two hours (AZ127) prior to analysis, the media were supplemented with 0.3% xylose to induce synthesis of the fusion proteins (+). Cells grown in the absence of xylose (-) are shown as controls.

**Figure 8–figure supplement 1.**
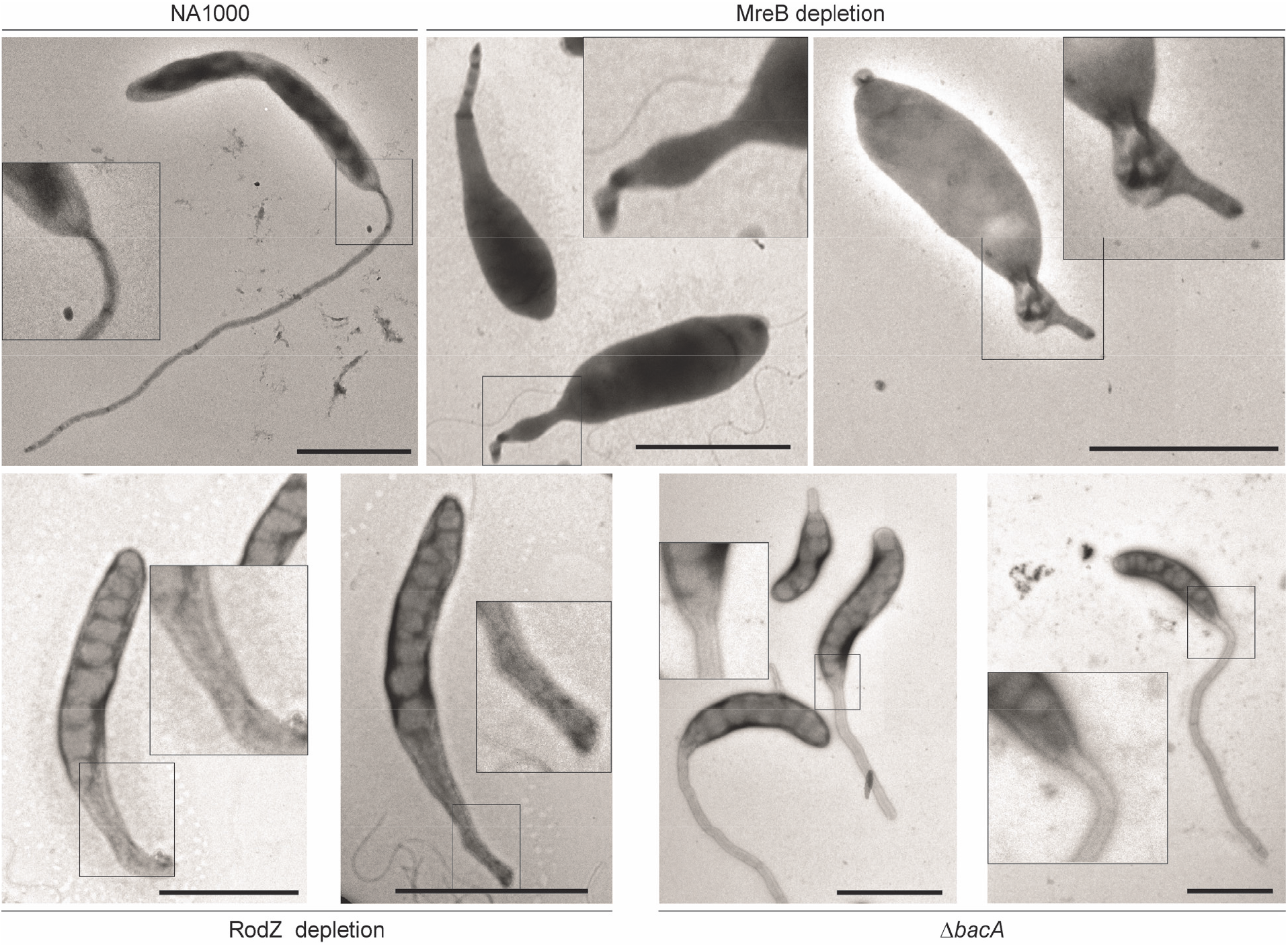
Transmission electron micrographs of mutants lacking scaffolding proteins. Shown are cells of strains NA1000 (WT), LS3809 (Δ*mreB* Pxyl::Pxyl-*mreB*), CJW2747 (Δ*rodZ::Ω* Pxyl::Pxyl-*rodZ*), and MT257 (Δ*bacA*) that were stained with uranyl acetate and visualized by transmission electron microscopy (scale bars: 2 μm). Strains NA1000 and MT257 (Δ*bacA*) was grown in M2G^-P^ for 24 h prior to imaging. Strains LS3809, and CJW2747 were grown to exponential phase in PYE medium containing the inducer xylose, washed, cultivated for 7 h in inducer-free PYE medium, and then diluted (1:20) into M2G^-P^ medium 24 h prior to imaging.

**Figure 12–figure supplement 1.**
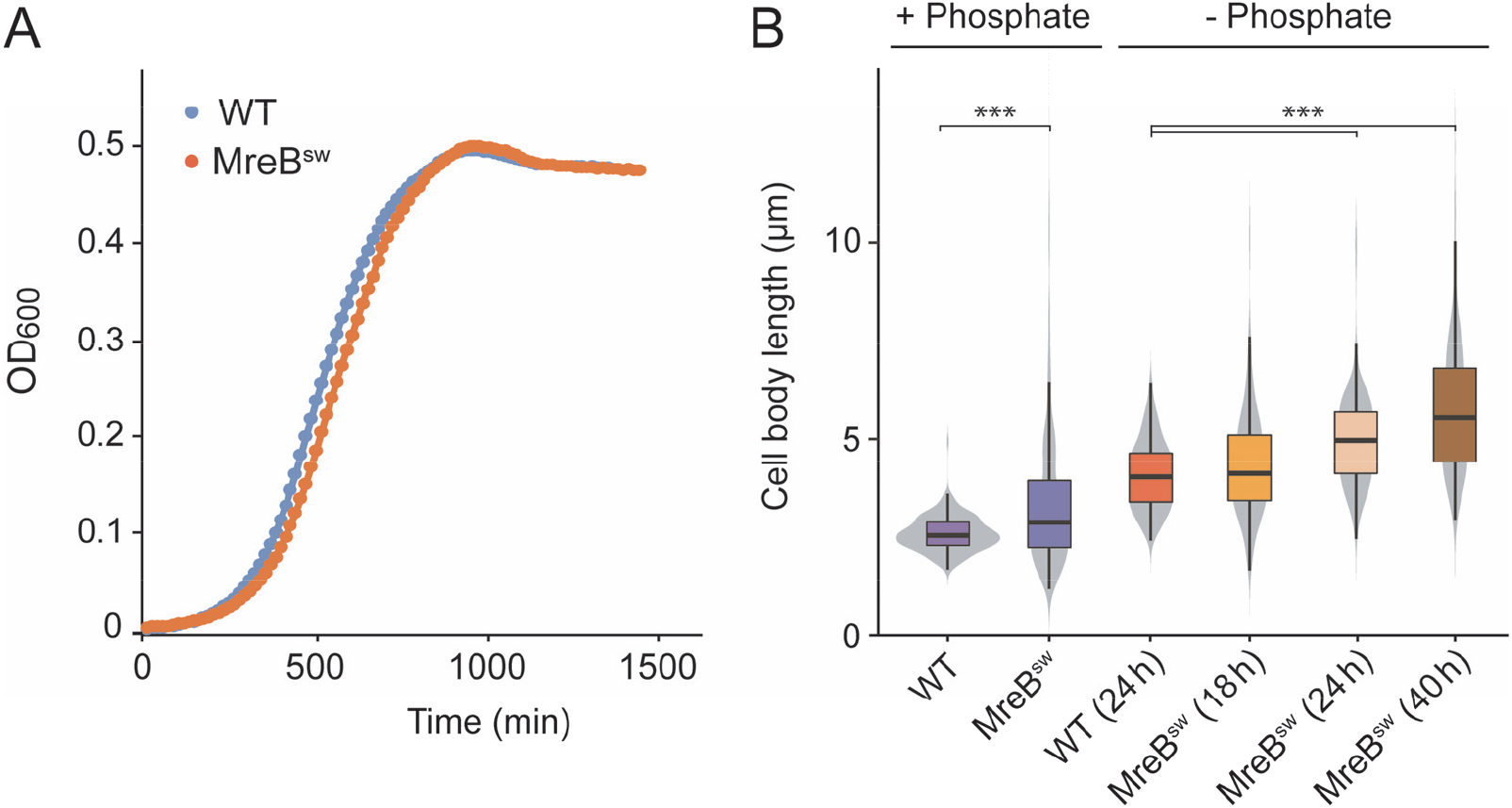
Growth characteristics of a strain producing an MreB-mCherry sandwich fusion. (**A**) Growth curves of strains NA1000 (WT) and MAB238 (mreB^sw^) in PYE medium. (**B**) Distribution of the cell body lengths in populations of strains NA1000 (WT) and MAB238 (*mreB^sw^*) during exponential growth in PYE medium or after cultivation for 18 h, 24 h and 40 h in M2G^-P^ medium. The values obtained (n=210 per strain) are shown as box plots, with the thick line indicating the median, the box the interquartile range and the wiskers the 2^nd^ and the 98^th^ percentile. In addition rotated kernel density plots (grey) are depicted for each dataset to indicate the distribution of the raw data (*** p < 10^-6^; t-test).

## Supplementary files

**Supplementary file 1.** Composition of cell body and stalk peptidoglycan from crossband- and LD-transpeptidase-deficient cells. The indicated strains were analyzed after growth for 24 h in M2G^-P^. Values are the mean ± variance of two independent experiments.

| Muropeptide | Relative percentage (%) in strain |  |  |  |  |  |
| --- | --- | --- | --- | --- | --- | --- |
| | WT (CB15N) | | ΔstpAB | | ΔldtDX | |
|  | Cell body | Stalk | Cell body | Stalk | Cell body | Stalk |
| Tri | 0.8±0.0 | 1.4±0.0 | 1.5±0.0 | 3.3±0.0 | n.d. | n.d. |
| Tetra | 16.5±0.4 | 11.1±6.6 | 19.3±0.3 | 12.1±3.5 | 19.3±0.1 | 13.6±0.0 |
| Penta | 7.9±0.0 | 6.5±0.1 | 8.9±0.0 | 6.6±0.0 | 8.6±0.0 | 7.8±0.0 |
| TriTri (LD) | 0.3±0.0 | 0.9±0.0 | 0.2±0.1 | 1.2±0.2 | n.d. | n.d. |
| TetraTri (LD ) | 1.0±0.1 | 2.9±0.1 | 0.6±0.0 | 2.2±0.2 | n.d. | n.d. |
| TriAnh/TetraTri (LD) | 1.1±0.0 | 1.3±0.0 | 1.1±0.0 | 1.9±0.0 | n.d. | n.d. |
| TetraTetra(LD) | 1.3±1.1 | 1.5±0.0 | 0.4±0.0 | 1.4±0.0 | 0.2±0.0 | 0.2±0.0 |
| TetraTetra | 12.7±0.0 | 10.8±1.5 | 14.6±0.1 | 10.4±0.1 | 14.3±0.0 | 13.5±0.0 |
| TetraPenta | 6.8±0.1 | 7.0±0.0 | 7.9±0.0 | 6.9±0.0 | 7.4±0.0 | 8.5±0.0 |
| TetraTetraTri or TetraTetraTri (LD) | 0.3±0.0 | 1.2±0.0 | 0.2±0.0 | 1.0±0.0 | n.d. | n.d. |
| TetraAnh | 1.4±0.0 | 1.7±0.0 | 1.1±0.0 | 0.8±0.0 | 1.2±0.0 | 1.2±0.0 |
| TetraTetraTetra | 3.8±0.0 | 5.1±0.0 | 4.1±0.0 | 4.8±0.1 | 4.2±0.0 | 5.9±0.0 |
| TetraTetraPenta | 1.8±0.0 | 3.0±0.0 | 1.9±0.0 | 2.6±0.0 | 2.1±0.0 | 3.2±0.0 |
| TetraTetraTetraTetra | n.d. | 0.3±0.2 | n.d. | 0.5±0.0 | n.d. | n.d. |
| TetraTetraTetraPenta | n.d. | 0.6±0.2 | n.d. | 0.3±0.0 | n.d. | n.d. |
| PentaAnh | 0.8±0.0 | 1.6±0.2 | 0.7±0.0 | 1.4±0.0 | 0.8±0.0 | 1.7±0.0 |
| TetraTetraAnh | 1.0±0.0 | 2.0±0.0 | 0.7±0.2 | 1.1±0.0 | 0.8±0.0 | 0.6±0.0 |
| TetraPentaAnh | 3.6±0.0 | 3.1±0.0 | 2.3±0.0 | 1.9±0.0 | 2.5±0.0 | 2.6±0.0 |
| TetraTetraTriAnh or TetraTetraTri(LD)Anh | 0.6±0.0 | 1.3±0.0 | 0.7±0.0 | 1.8±0.0 | 1.3±0.0 | 1.9±0.0 |
| TetraTetraTetraAnh | 6.4±0.1 | 6.2±0.0 | 6.8±0.0 | 6.2±0.0 | 6.8±0.0 | 6.8±0.0 |
| TetraTetraPentaAnh | 6.6±0.1 | 6.2±0.0 | 6.4±0.0 | 5.7±0.1 | 6.7±0.0 | 6.6±0.0 |
| TetraTetraTetraPentaAnh I | 1.4±0.0 | 1.0±0.0 | 1.3±0.0 | 1.1±0.2 | 1.4±0.0 | 2.0±0.0 |
| TetraTetraTetraPentaAnh II | 1.4±0.0 | 1.3±0.0 | 1.5±0.0 | 1.2±0.0 | 1.7±0.0 | 1.8±0.0 |
| TetraTetraTetradiAnh | 1.4±0.0 | 1.5±0.0 | 1.8±0.0 | 1.4±0.5 | 1.4±0.0 | 1.6±0.7 |
| TetraTetraTetraPentadiAnh | 1.8±0.3 | 1.3±0.0 | 1.9±0.2 | 1.1±0.0 | 2.3±0.0 | 1.6±0.3 |
| TetraTetradiAnh | 1.8±0.0 | 1.3±0.0 | 1.2±0.4 | 1.0±0.0 | 2.7±0.0 | 1.6±0.0 |
| TetraTetraPentadiAnh | 0.7±0.0 | 0.4±0.0 | 0.7±0.0 | 0.4±0.0 | 0.9±0.0 | 0.8±0.0 |
| all known | 83.3±0.6 | 82.5±4.7 | 87.7±0.5 | 80.3±0.0 | 86.5±0.0 | 83.4±0.2 |
| Monomers (total) | 33.0±0.1 | 27.0±4.5 | 36.0±1.1 | 30.1±4.3 | 34.6±0.3 | 29.0±0.4 |
| Dimers (total) | 35.6±3.4 | 37.2±0.0 | 32.9±0.7 | 34.8±1.6 | 32.2±0.0 | 32.3±0.0 |
| Trimers (total) | 26.0±0.5 | 30.2±3.7 | 25.9±0.1 | 29.9±4.1 | 27.0±0.1 | 32.2±1.7 |
| Tetramers (total) | 5.4±0.6 | 5.5±0.2 | 5.3±0.2 | 5.2±1.4 | 6.2±0.0 | 6.5±0.4 |
| Tripeptides (total) | 3.0±0.0 | 6.4± 0.6 | 3.1±0.0 | 9.3±1.3 | 0.5±0.0 | 0.8±0.0 |
| Tetrapeptides (total) | 74.7±0.0 | 71.9±2.5 | 74.5±0.3 | 69.2±1.3 | 77.6±0.0 | 75.4±0.2 |
| Pentapeptides (total) | 21.7±0.0 | 21.1±0.5 | 21.6±0.0 | 20.2±0.0 | 21.5±0.0 | 23.2±0.2 |
| 3-3 crosslinks | 2.3±0.6 | 5.2±0.7 | 1.0±0.0 | 4.2±0.6 | 0.6±0.0 | 0.9±0.0 |
| Chain ends (Anh) | 16.5±0.2 | 17.2±0.9 | 14.2±1.2 | 14.5±0.8 | 17.0±0.0 | 17.4±0.3 |
| Degree of cross-linkage<sup>1</sup> | 39.2±0.0 | 42.9±2.2 | 37.6±0.4 | 41.2±1.2 | 38.8±0.1 | 42.5±0.2 |
| % peptides in cross-links | 67.0±0.1 | 73.0±4.5 | 64.0±1.1 | 69.9±4.3 | 65.4±0.3 | 71.0±0.4 |
| Mean of disaccharides units ± SD (n=2) |  |  |  |  |  |  |
| Average glycan chain length | 6.0±0.1 | 5.8±0.1 | 7.1±0.3 | 6.9±0.2 | 5.9±0.0 | 5.8±0.0 |

**Supplementary file 2.**
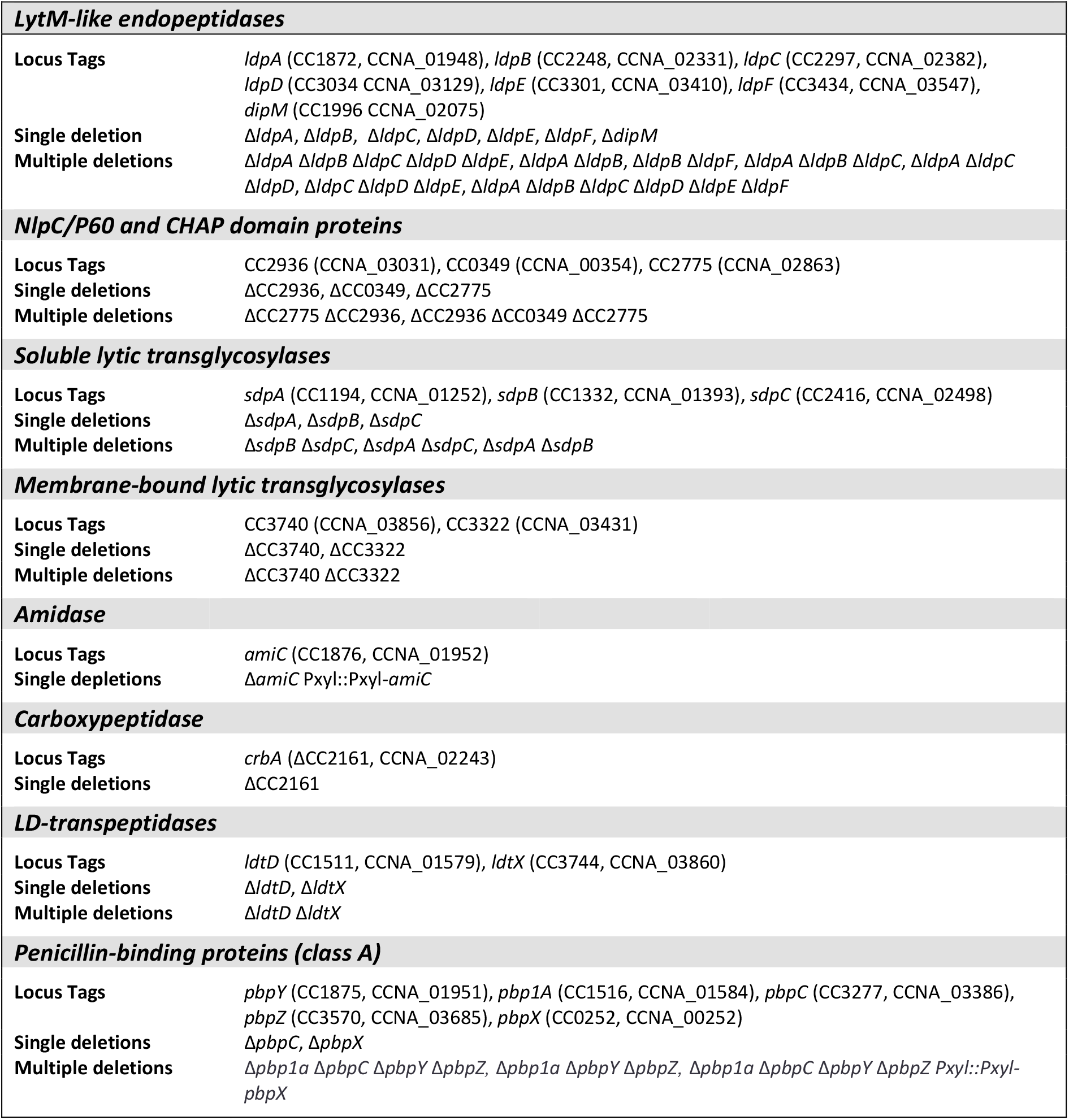
List of mutant strains analyzed for stalk defects.

**Supplementary file 3.** Composition of peptidoglycan isolated from autolysin-deficient cells. The indicated strains were analyzed after growth for 24 h in M2G^-P^. The values are the mean ± variance of two independent experiments.

| Muropeptide | Relative percentage (%) in strain |  |  |  |  |
| --- | --- | --- | --- | --- | --- |
| | WT | $\Delta dipM$ | $\Delta crbA$ | $\Delta ldpA$ | $\Delta sd pAB$ |
| Tri | 0.9±0.0 | 1.8±0.1 | 0.6±0.0 | 1.1±0.0 | 4.3±2.1 |
| Tetra | 17.5±0.0 | 21.3±0.5 | 9.8±0.5 | 18.2±0.1 | 27.5±1.8 |
| Penta | 7.9±0.1 | 8.1±0.3 | 11.0±0.0 | 7.4±0.0 | 6.6±0.0 |
| TriTri (LD) | 0.2±0.0 | 0.2±0.0 | 0.2±0.0 | 0.2±0.0 | 0.5±0.0 |
| TetraTri (LD) | 1.1±0.0 | 0.9±0.0 | 0.2±0.0 | 0.8±0.0 | 1.9±0.0 |
| TriAnh +small amount of TetraTri (LD) | 0.9±0.0 | 1.0±0.0 | 0.5±0.0 | 0.9±0.0 | 1.6±0.3 |
| TetraTetra (LD) | 0.2±0.0 | 0.5±0.0 | 1.1±0.0 | 0.9±0.5 | 0.6±0.0 |
| TetraTetra | 15.1±0.0 | 15.9±0.0 | 10.8±0.3 | 15.8±0.0 | 16.7±0.8 |
| TetraPenta | 7.3±0.2 | 7.3±0.1 | 12.1±0.1 | 7.1±0.1 | 5.3±0.0 |
| TetraTetraTri or TetraTetraTri (LD) | 0.4±0.0 | 0.3±0.0 | 0.3±0.0 | 0.3±0.0 | 0.4±0.0 |
| TetraAnh | 0.3±0.0 | 0.5±0.0 | 0.2±0.0 | 0.5±0.0 | 0.8±0.0 |
| TetraTetraTetra | 4.4±0.1 | 4.5±0.0 | 3.5±0.2 | 4.6±0.0 | 5.5±0.2 |
| TetraTetraPenta | 1.8±0.0 | 1.8±0.0 | 3.6±0.0 | 1.8±0.0 | 1.5±0.0 |
| TetraTetraTetraTetra | 0.8±0.0 | 0.8±0.0 | 0.6±0.0 | 0.8±0.0 | 1.1±0.0 |
| TetraTetraTetraPenta | 0.4±0.0 | 0.4±0.0 | 0.6±0.0 | 0.4±0.0 | 0.4±0.0 |
| PentaAnh | 0.8±0.0 | 0.8±0.0 | 0.8±0.0 | 0.8±0.0 | 1.7±0.0 |
| TetraTetraAnh | 4.3±0.0 | 3.9±0.0 | 3.0±0.0 | 4.1±0.0 | 2.6±0.0 |
| TetraPentaAnh | 0.9±0.0 | 0.8±0.0 | 0.4±0.1 | 0.6±0.0 | 0.9±0.1 |
| TetraTetraTriAnh or TetraTetraTri(LD)Anh | n.d. | n.d. | n.d. | n.d. | n.d. |
| TetraTetraTetraAnh | 7.6±0.1 | 7.1±0.0 | 5.6±0.0 | 7.6±0.0 | 6.4±0.0 |
| TetraTetraPentaAnh | 6.4±0.5 | 5.9±0.1 | 7.0±0.0 | 7.0±0.0 | 4.0±0.0 |
| TetraTetraTetraPentaAnh I | 1.2±0.0 | 1.2±0.0 | 1.3±0.0 | 1.3±0.0 | 1.0±0.0 |
| TetraTetraTetraPentaAnh II | 1.3±0.0 | 1.3±0.0 | 2.3±0.0 | 1.4±0.0 | 0.7±0.0 |
| TetraTetraTetradiAnh | 1.1±0.0 | 1.0±0.0 | 1.1±0.0 | 1.2±0.0 | 0.5±0.0 |
| TetraTetraTetraPentadiAnh | 0.9±0.0 | 0.9±0.0 | 1.3±0.0 | 1.0±0.0 | 0.3±0.0 |
| TetraTetradiAnh | 1.5±0.0 | 1.4±0.0 | 1.8±0.0 | 1.5±0.0 | 0.4±0.0 |
| TetraTetraPentadiAnh | 0.7±0.0 | 0.6±0.0 | 1.2±0.0 | 0.7±0.0 | 0.3±0.0 |
| all known | 85.7±1.7 | 90.1±1.0 | 80.6±0.0 | 87.9±0.1 | 93.4±0.5 |
| Monomers (total) | 32.9±0.5 | 37.3±2.1 | 28.4±0.2 | 32.9±0.1 | 45.4±1.5 |
| Dimers (total) | 35.7±1.7 | 34.1±0.0 | 36.5±1.4 | 35.2±1.1 | 31.0±0.2 |
| Trimers (total) | 26.0±1.5 | 23.6±1.1 | 27.6±1.3 | 26.5±0.1 | 20.0±0.4 |
| Tetramers (total) | 5.4±0.6 | 5.1±0.3 | 7.4±0.2 | 5.5±0.3 | 3.7±0.0 |
| Tripeptides (total) | 3.1±0.0 | 4.0±0.2 | 1.8±0.3 | 3.0±0.0 | 8.1±5.4 |
| Tetrapeptides (total) | 77.1±0.2 | 77.0±0.6 | 68.9±0.6 | 77.9±0.0 | 76.3±6.9 |
| Pentapeptides (total) | 19.5±0.1 | 18.5±0.2 | 29.0±0.0 | 18.5±0.1 | 14.8±0.0 |
| 3-3 crosslinks | 1.0±0.0 | 1.0±0.0 | 1.0±0.0 | 1.2±0.2 | 1.8±0.0 |
| Chain ends (Anh) | 15.0±0.2 | 13.9±0.1 | 15.2±0.4 | 15.1±0.0 | 11.6±1.0 |
| Degree of cross-linkage<sup>1</sup> | 39.2±0.6 | 36.6±1.1 | 42.2±0.2 | 39.3±0.0 | 31.6±0.5 |
| % Peptides in crosslinks | 67.1±0.5 | 62.7±2.1 | 71.6±0.2 | 67.1±0.1 | 54.6±1.5 |
| Mean of disaccharides units ± SD (n=2) |  |  |  |  |  |
| Average glycan chain length | 6.7±0.0 | 7.2±0.0 | 6.6±0.1 | 6.6±0.0 | 8.6±0.6 |

**Supplementary file 4.** Composition of peptidoglycan isolated from cells with defects in scaffolding proteins. Strains MT257 (Δ*bacA*), MAB238 (*mreB::mreB^sw^*), and NA1000 (WT) were grown for 24 h in M2G^-P^ prior to analysis. Strains CJW2747 (Δ*rodZ::Ω* Pxyl::Pxyl-*rodZ*) and LS3809 (Δ*mreB* Pxyl::Pxyl-*mreB*) were first cultivated for 7 h in PYE and then grown for 24 h in M2G^-P^. Values are the mean values ± variance of two independent experiments.

| Muropeptide | Relative percentage (%) in strain |  |  |  |  |
| --- | --- | --- | --- | --- | --- |
| | WT | $\Delta bacA$ | RodZ depl. | MreB depl. | MreB<sup>SW</sup> |
| Tri | 0.9±0.0 | 1.4±0.0 | 5.0±0.0 | 5.1±0.1 | 3.8±0.0 |
| Tetra | 17.5±0.0 | 19.6±0.0 | 17.7±0.2 | 17.4±0.0 | 19.2±0.1 |
| Penta | 7.9±0.1 | 8.0±0.1 | 8.3±0.0 | 8.7±0.0 | 7.3±0.0 |
| TriTri (LD) | 0.2±0.0 | 0.2±0.0 | 0.8±0.0 | 1.0±0.0 | 0.5±0.0 |
| TetraTri (LD) | 1.1±0.0 | 0.7±0.0 | 1.9±0.0 | 2.5±0.0 | 1.9±0.0 |
| TriAnh/TetraTri (LD) | 0.9±0.0 | 1.1±0.0 | 3.6±0.0 | 3.7±0.0 | 3.0±0.0 |
| TetraTetra (LD) | 0.2±0.0 | 0.4±0.0 | 1.1±0.0 | 1.3±0.0 | 0.9±0.0 |
| TetraTetra | 15.1±0.0 | 16.8±0.2 | 13.6±0.0 | 13.3±0.1 | 15.8±0.0 |
| TetraPenta | 7.3±0.2 | 7.7±0.0 | 6.4±0.0 | 6.8±0.0 | 5.9±0.0 |
| TetraTetraTri or TetraTetraTri (LD) | 0.4±0.0 | 0.3±0.0 | 0.6±0.0 | 0.7±0.0 | 0.6±0.0 |
| TetraAnh | 0.3±0.0 | 0.5±0.0 | 0.9±0.0 | 0.8±0.0 | 0.8±0.0 |
| TetraTetraTetra | 4.4±0.1 | 5.0±0.0 | 3.7±0.0 | 3.1±0.0 | 4.4±0.0 |
| TetraTetraPenta | 1.8±0.0 | 1.9±0.0 | 1.4±0.0 | 1.3±0.0 | 1.3±0.0 |
| TetraTetraTetraTetra | 0.8±0.0 | 0.9±0.0 | 0.3±0.0 | 0.3±0.0 | 0.2±0.0 |
| TetraTetraTetraPenta | 0.4±0.0 | 0.4±0.0 | 0.1±0.0 | 0.2±0.0 | 0.1±0.0 |
| PentaAnh | 0.8±0.0 | 0.9±0.0 | 0.6±0.0 | 0.5±0.0 | 0.7±0.0 |
| TetraTetraAnh | 4.3±0.0 | 4.2±0.0 | 0.7±0.0 | 0.7±0.0 | 0.6±0.0 |
| TetraPentaAnh | 0.9±0.0 | 0.8±0.0 | 2.2±0.0 | 2.2±0.0 | 2.2±0.0 |
| TetraTetraTriAnh or TetraTetraTri(LD)Anh | n.d. | n.d. | 1.5±0.0 | 1.5±0.0 | 1.4±0.0 |
| TetraTetraTetraAnh | 7.6±0.1 | 7.0±0.0 | 5.5±0.0 | 4.9±0.1 | 6.1±0.0 |
| TetraTetraPentaAnh | 6.4±0.5 | 6.1±0.0 | 4.3±0.1 | 4.1±0.0 | 4.7±0.0 |
| TetraTetraTetraPentaAnh I | 1.2±0.0 | 1.3±0.0 | 0.6±0.0 | 0.4±0.0 | 0.7±0.0 |
| TetraTetraTetraPentaAnh II | 1.3±0.0 | 1.3±0.0 | 0.6±0.0 | 0.5±0.0 | 0.7±0.0 |
| TetraTetraTetradiAnh | 1.1±0.0 | 1.0±0.0 | 1.0±0.0 | 0.7±0.0 | 0.9±0.0 |
| TetraTetraTetraPentadiAnh | 0.9±0.0 | 0.9±0.0 | 1.3±0.0 | 1.0±0.0 | 1.3±0.0 |
| TetraTetradiAnh | 1.5±0.0 | 1.2±0.0 | 1.3±0.0 | 1.1±0.0 | 1.2±0.0 |
| TetraTetraPentadiAnh | 0.7±0.0 | 0.6±0.0 | 0.4±0.0 | 0.4±0.0 | 0.3±0.0 |
| all known | 85.7±1.7 | 90.2±0.0 | 85.3±0.1 | 84.4±0.2 | 86.4±0.1 |
| Monomers (total) | 32.9±0.5 | 35.1±0.0 | 38.2±0.6 | 38.6±0.3 | 36.8±0.2 |
| Dimers (total) | 35.7±1.7 | 35.4±0.0 | 36.9±0.1 | 38.7±0.0 | 36.8±0.0 |
| Trimers (total) | 26.0±1.5 | 24.3±0.0 | 21.5±0.2 | 19.7±0.1 | 22.8±0.1 |
| Tetramers (total) | 5.4±0.6 | 5.2±0.0 | 3.4±0.0 | 2.9±0.0 | 3.5±0.0 |
| Tripeptides (total) | 3.1±0.0 | 3.5±0.0 | 10.8±0.0 | 11.8±0.4 | 8.5±0.1 |
| Tetrapeptides (total) | 77.1±0.2 | 77.0±0.0 | 70.6±0.0 | 69.1±0.6 | 74.3±0.0 |
| Pentapeptides (total) | 19.5±0.1 | 18.9±0.0 | 18.6±0.0 | 19.1±0.0 | 17.2±0.0 |
| 3-3 crosslinks | 1.0±0.0 | 0.8±0.0 | 3.0±0.0 | 3.7±0.0 | 2.6±0.0 |
| Chain ends (Anh) | 15.0±0.2 | 14.1±0.0 | 11.3±0.0 | 10.4±0.0 | 11.4±0.1 |
| Degree of cross-linkage<sup>1</sup> | 39.2±0.6 | 37.8±0.0 | 35.3±0.2 | 34.7±0.1 | 36.3±0.1 |
| % peptides in cross-links | 67.1±0.5 | 64.9±0.0 | 61.8±0.6 | 61.4±0.3 | 63.2±0.2 |
| Mean of disaccharides units ± standard deviation (n=2) | | | | | |
| Average glycan chain length | 6.7±0.0 | 7.1±0.0 | 8.8±0.0 | 9.6±0.0 | 8.8±0.0 |

**Supplementary file 5.**
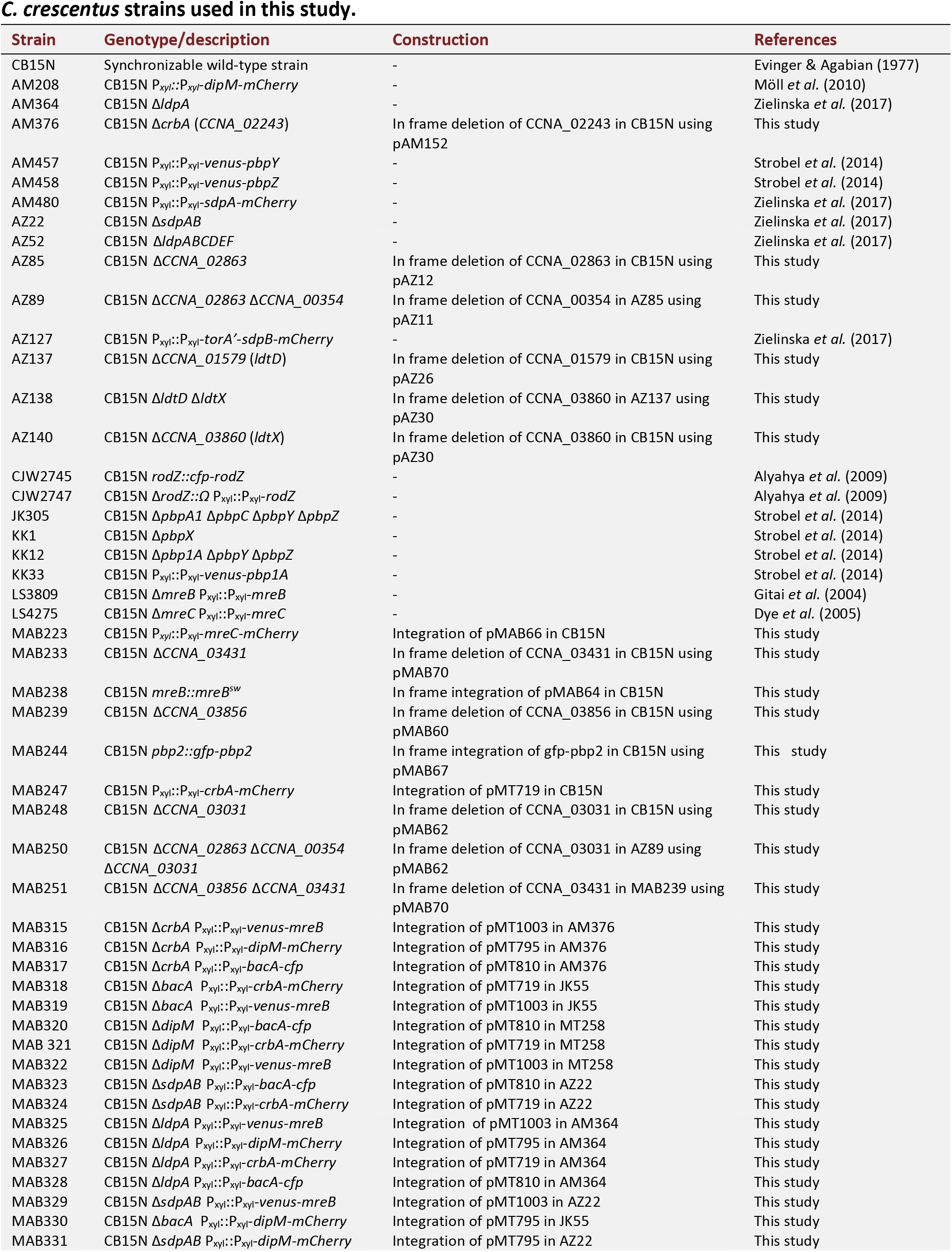
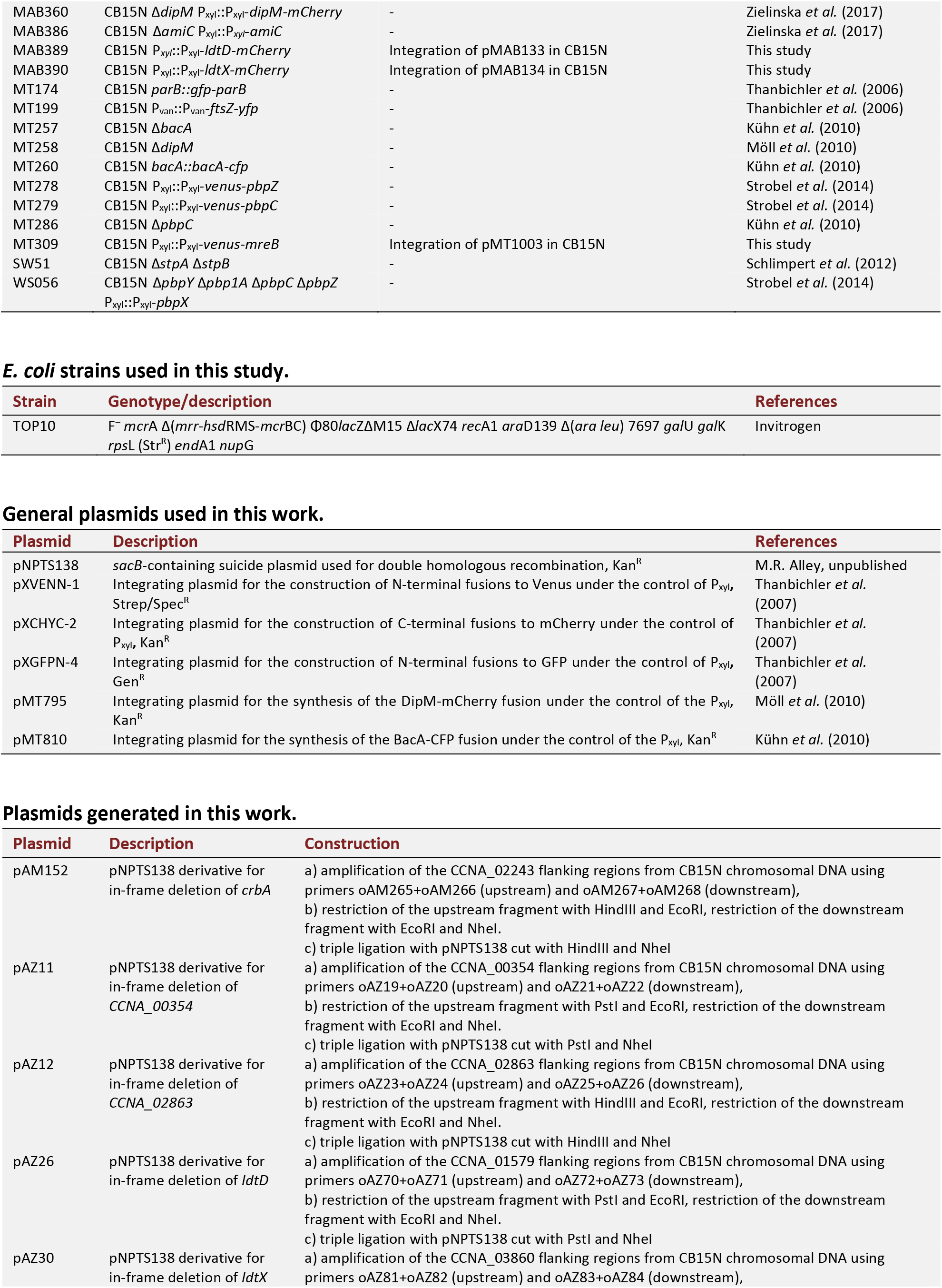
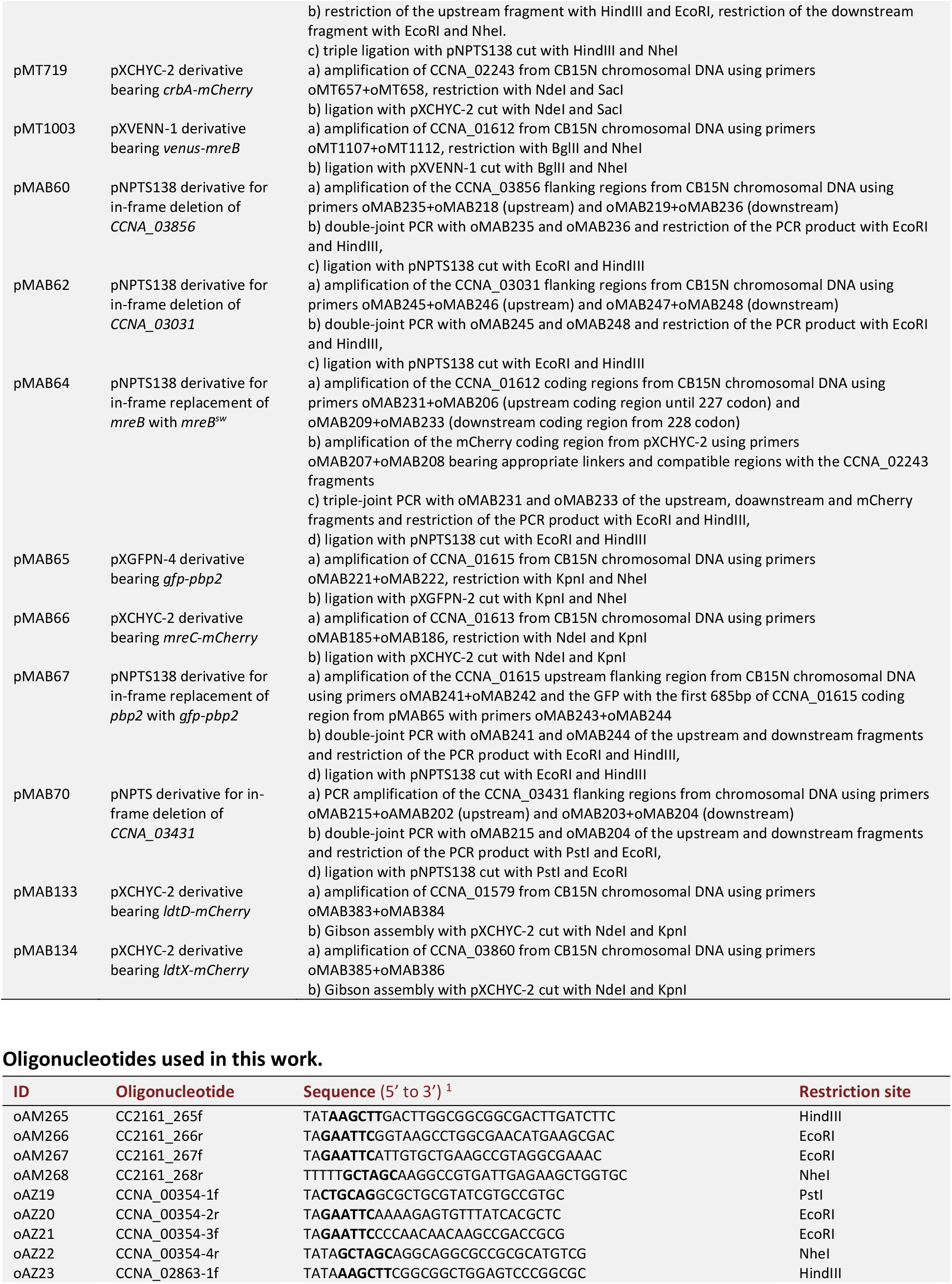
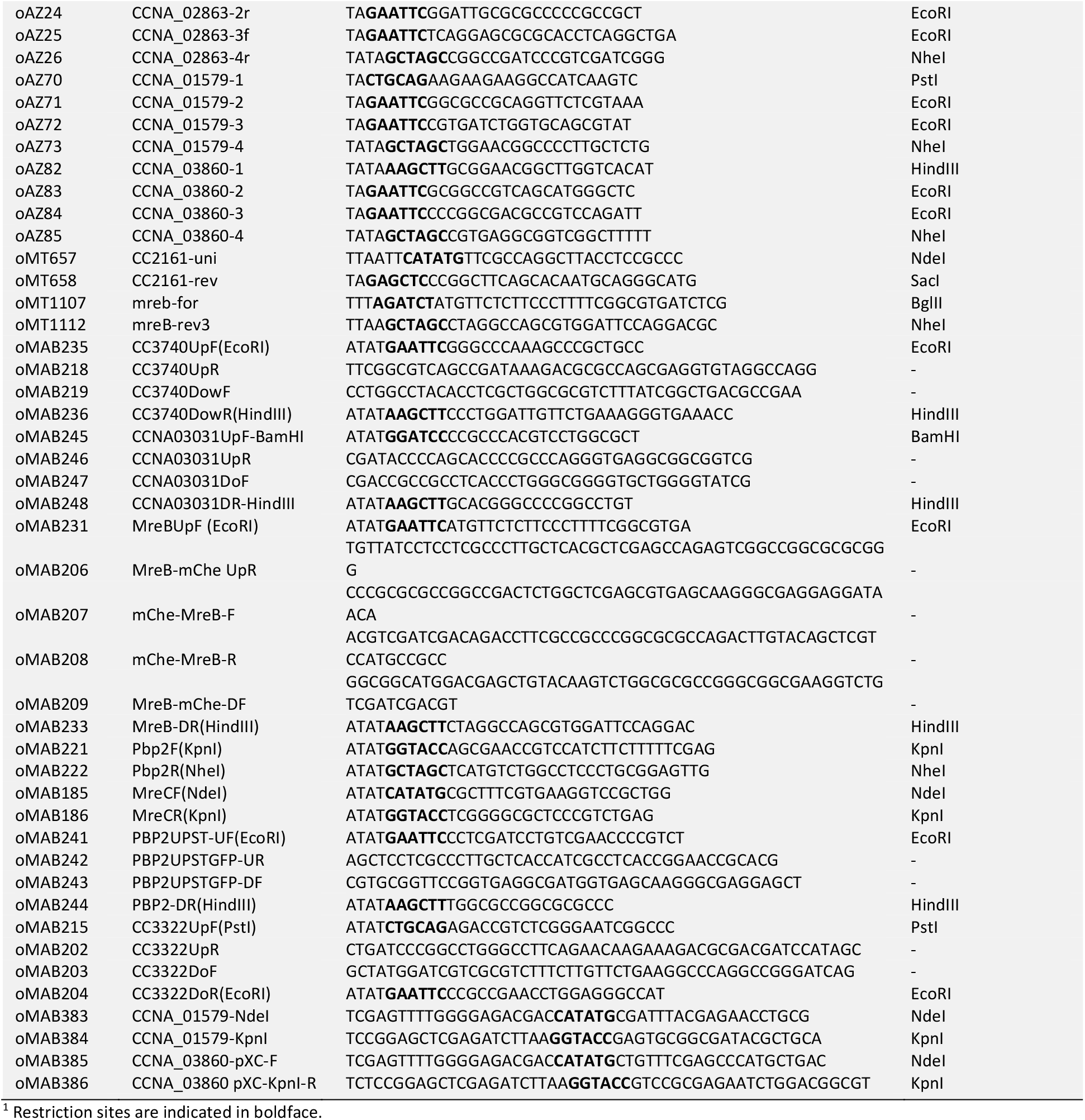
Strains, plasmids and oligonucleotides.

